# The Structural Logic of Dynamic Signaling in the *Escherichia coli* Serine Chemoreceptor

**DOI:** 10.1101/2024.07.23.604838

**Authors:** Georgina I. Reyes, Caralyn E. Flack, John S. Parkinson

## Abstract

The experimental challenges posed by integral membrane proteins hinder molecular understanding of transmembrane signaling mechanisms. Here, we exploited protein crosslinking assays in living cells to follow conformational and dynamic stimulus signals in Tsr, the *Escherichia coli* serine chemoreceptor. Tsr mediates serine chemotaxis by integrating transmembrane serine-binding inputs with adaptational modifications of a methylation helix bundle to regulate a signaling kinase at the cytoplasmic tip of the receptor molecule. We created a series of cysteine replacements at Tsr residues adjacent to hydrophobic packing faces of the bundle helices and crosslinked them with a cell-permeable, bifunctional thiol-reagent. We identified an extensively crosslinked dynamic junction midway through the methylation helix bundle that seemed uniquely poised to respond to serine signals. We explored its role in mediating signaling shifts between different packing arrangements of the bundle helices by measuring crosslinking in receptor molecules with apposed pairs of cysteine reporters in each subunit and assessing their signaling behaviors with an in vivo kinase assay. In the absence of serine, the bundle helices evinced compact kinase-ON packing arrangements; in the presence of serine, the dynamic junction destabilized adjacent bundle segments and shifted the bundle to an expanded, less stable kinase-OFF helix-packing arrangement. An AlphaFold 3 model of kinase-active Tsr showed a prominent bulge and kink at the dynamic junction that might antagonize stable structure at the receptor tip. Serine stimuli probably inhibit kinase activity by shifting the bundle to a less stably-packed conformation that relaxes structural strain at the receptor tip, thereby freezing kinase activity.

**SIGNIFICANCE:** This study used in vivo protein crosslinking to follow stimulus-induced signals through Tsr, a bacterial transmembrane receptor for chemotaxis to serine. Our experiments distinguished Tsr conformations that activate or inhibit its signaling kinase partner and showed how those signals reach the receptor’s kinase-controlling cytoplasmic tip. A dynamic junction in the Tsr molecule triggers stimulus responses by propagating a less stable helix-packing arrangement to flanking structural elements, thereby reducing structural stresses at the receptor tip. An AlphaFold model indicated that the dynamic junction might cause a structural distortion that destabilizes the receptor tip in the absence of a serine stimulus. This model and experimental approach could help to elucidate the signaling logic and mechanisms in other transmembrane chemoreceptor proteins.

## INTRODUCTION

The chemotaxis machinery of *Escherichia coli* has long provided a powerful experimental system for investigating molecular mechanisms of stimulus detection and signaling by transmembrane chemoreceptors (1–3). The best understood bacterial chemoreceptors belong to the methyl-accepting chemotaxis protein (MCP) superfamily (4). MCP molecules are homodimeric, mainly alpha-helical proteins that assemble signaling complexes at their cytoplasmic hairpin tips through interactions with two soluble partner proteins, a histidine autokinase (CheA) and a scaffolding protein (CheW) that couples CheA activity to chemoreceptor control (Fig. 1*A*; Fig. S1). The fundamental unit of chemoreceptor activity is a core signaling unit comprising six receptor molecules organized as two trimers of dimers, one CheA homodimer, and two CheW protomers (5) (Fig. S1). These signaling complexes are in turn networked into large cooperative signaling arrays through hexameric CheA-CheW and CheW-CheW rings (6, 7).

**Fig. 1.**
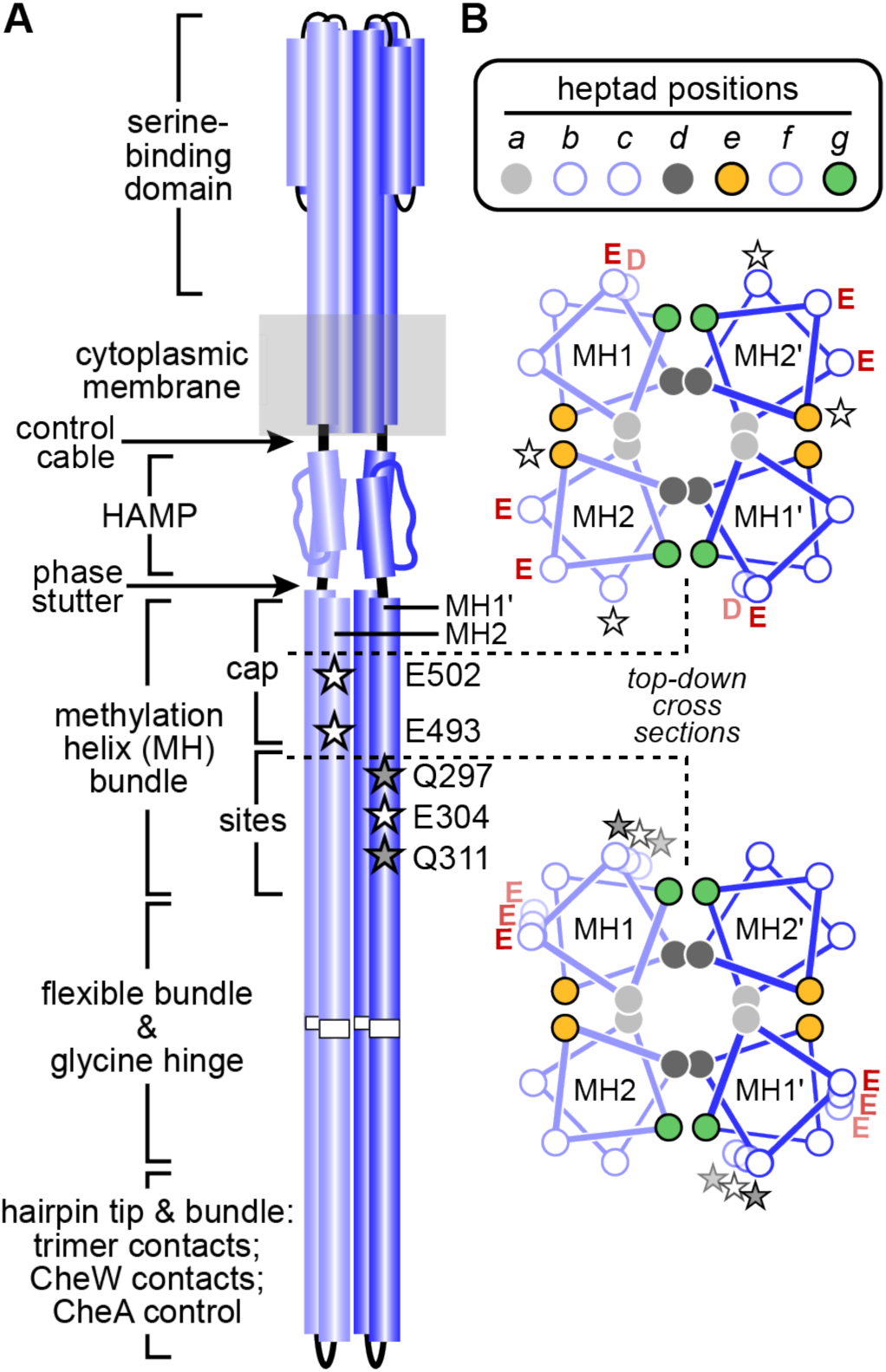
Tsr structural features. (A) Signaling elements in the Tsr homodimer. Cylinders represent alpha-helical segments of Tsr subunits (light and dark blue), drawn approximately to scale. The cytoplasmic segment below the HAMP domain is an antiparallel, 4-helix bundle. Stars indicate adaptational modification site residues, E (white) and Q (gray). (B) The cap and sites segments of the MH bundle. Helical wheels depict cross-sections (viewed in the top-down, HAMP to tip direction) of a four-helix knobs-in-holes (*a-d*) packing configuration. The descending helices (MH1, MH1’) align with the ascending helices (MH2, MH2’) in anti-parallel orientation. Each helix contains a series of seven-residue (heptad) repeats that comprise two helical turns and packing layers. Hydrophobic residues at *a* and *d* heptad positions promote the principal packing interactions between the bundle helices; edge residues at *e* and *g* heptad positions play ancillary roles in bundle geometry and stability. Adaptation site residues (stars) and adjacent D or E residues lie at solvent-exposed positions (*b*, *c, f*) that create extended acidic faces that modulate helix stability and packing in the MH bundle.

Much of what we know about the molecular mechanisms of MCP signaling has come from studies of the *E. coli* Tsr (serine) and Tar (aspartate) chemoreceptors. In isotropic chemical environments, Tsr and activate their CheA partners, which autophosphorylate using ATP, then donate their phosphoryl groups to the CheY response regulator [reviewed in (2)]. Phospho-CheY molecules interact with flagellar basal bodies to initiate directional changes that produce a random-walk swimming pattern. Upon sensing an attractant increase, Tsr and Tar inhibit CheA, halting the flux of phosphoryl groups to CheY to promote forward swimming, the default motor behavior. Cellular phospho-CheY is short-lived due to the action of a dedicated phosphatase (CheZ), thereby ensuring rapid behavioral responses to chemoeffector stimuli.

A sensory adaptation system modulates the ligand sensitivity of the Tsr and Tar kinase control responses, enabling swimming cells to detect spatial chemoeffector gradients in temporal fashion by comparing current chemical conditions with those averaged over the past few seconds of their travels [reviewed in (2)]. Two MCP-specific enzymes, CheR, a methyltransferase, and CheB, a methylesterase, respectively methylate or demethylate glutamyl residues in the cytoplasmic portion of the receptor molecule to adjust its signaling activity to prevailing chemoeffector levels. Under steady-state conditions, CheB demethylates receptors in the kinase-ON state and shifts them toward kinase-OFF output, whereas CheR methylates receptor molecules in the kinase-OFF state and shifts them toward kinase-ON output (2).

Adaptational modifications adjust ligand sensitivity of the Tsr and Tar kinase control responses over more than a hundred-fold range but produce only a few-fold shift in their ligand-binding affinities (8–13). This mechanistic mystery has remained unsolved for decades. How can the receptor’s mechanisms of ligand and sensory adaptation control of kinase activity operate so far out of equilibrium? To do so, ligand binding and sensory adaptation must modulate different structural properties of the signaling elements in receptor molecules (14–17).

The signaling architecture of MCP molecules can promote sensitive responses to small changes in chemoeffector concentration by coupling adjacent structural elements in dynamic opposition through special linkers (Fig. 1*A*). Ligand-binding information from the periplasmic sensing domain and 4-helix transmembrane bundle modulates the stability and/or packing arrangement of a 4-helix HAMP bundle (18, 19) through a five-residue control cable helix at the cytoplasmic side of the membrane (20–22) (Fig. 1*A*). (HAMP domains are versatile input-output relays in many microbial signaling proteins, particularly sensor <u>H</u>istidine kinases, <u>A</u>denylyl cyclases, <u>M</u>CPs and some <u>P</u>hosphatases) [reviewed in (23)]. The HAMP domain, through a 4-residue “phase stutter” connection, in turn influences the packing stability or geometry of the methylation helix (MH) bundle (16, 19, 24) (Fig. 1*A*). Structural signals from the MH bundle travel to the hairpin bundle and tip through an intervening flexible bundle and glycine hinge (25–27) that may couple their signaling structures in dynamic opposition as well (28).

The dynamic behaviors of the MH bundle have been extensively characterized with in vitro studies of the aspartate receptor Tar (14, 28–34, 35, 36–41). Those studies have shown that the N-terminal MH1 helix is more dynamic than the C-terminal MH2 helix (37) over nanosecond (38) to millisecond (42) timescales. The dynamic motions of both helices involve fluctuations in helicity and backbone motions, presumably reflecting transient excursions from stable helix-packing interactions in the four-helix bundle (35, 37, 38, 43). Adaptational modifications influence the dynamic behaviors of both helices (14, 40), at least in part by altering the density of helix-destabilizing negative charges on solvent-exposed acidic faces of both helices (33) (Fig. 1*B*). Overall, in Tar the MH1 helix is more dynamic than its MH2 partner and their packing interactions respond to adaptational modifications (14, 35, 42) and to structural changes induced by attractant ligands (13, 41) or by kinase-OFF receptor lesions (38).

In the present study we explored stimulus-triggered conformational and dynamic changes in the MH bundle of the serine receptor Tsr. We tracked these events in living cells through in vivo protein crosslinking methods that elucidate structural features of chemoreceptors (16, 44–46) and their higher-order signaling complexes (47–50) under conditions that are difficult to replicate faithfully in vitro. Combined with in vivo FRET kinase assays, receptor crosslinking provides functional snapshots of helix packing configurations and motions for different signaling states in native receptor arrays (16). With these approaches we have been able to follow the transmission of stimulus-induced signals through Tsr in unprecedented detail and have identified structural features that collectively regulate signal transmission and kinase output state.

## RESULTS

### Scanning for Dynamic Sites in the Tsr MH Bundle

Helix packing in coiled-coils occurs mainly through hydrophobic interactions between residues at *a* and *d* positions in a repeating heptad pattern (51) (Fig. 1*B*). Residues at *e* and *g* heptad positions flank the hydrophobic core residues and can contribute to packing stability (51), but are less critical to receptor function than are the hydrophobic packing residues at *a* and *d* heptad positions (16, 24, 29, 34). To survey in vivo dynamic behaviors of the MH bundle helices we constructed mutations in Tsr expression plasmid pRR53 (47) that introduced a cysteine replacement at an *e* or *g* heptad position. In all, we created 30 mutant plasmids, each encoding Tsr subunits with one cysteine residue (single-CYS). When expressed in a receptorless but otherwise wild-type host (UU2612), 24/30 of the mutant plasmids supported demonstrable serine chemotaxis in soft agar assays (Table S1) and all reporter proteins had intracellular amounts within two-fold of the wild-type level (Table S1), consistent with a native or near-native structure.

We expressed single-CYS receptors from the mutant plasmids in receptorless strain UU2610, which lacked the sensory adaptation enzymes CheR and CheB to simplify receptor immunoblot patterns (16). Importantly, this host contained wild-type CheA and CheW to allow the receptors to assemble core signaling units (CSUs) (Fig. S1) and arrays. We induced crosslinking by treating the cells with bismaleimidoethane (BMOE), a cell-permeable, bifunctional thiol-reactive crosslinking reagent with an 8 Å spacer (6, 16, 52), using the same reaction conditions for all reporters (200 µM BMOE, 100”, 30°C). Cell lysates were subjected to denaturing sodium dodecyl sulfate-polyacrylamide gel electrophoresis (SDS-PAGE) and Tsr subunits were detected and quantified by immunoblotting with a polyclonal rabbit antiserum raised against the highly conserved Tsr hairpin tip (48, 53). Fig. S2*A* shows examples of the SDS-PAGE immunoblots.

We expected that dynamic motions of the MH bundle helices might bring the cysteine reporters in a Tsr dimer within BMOE crosslinking distance (5-10 Å) (Fig. S1). It seemed less likely that BMOE would promote crosslinks between the receptor molecules in adjacent dimers within core signaling units because of the relatively long distances between the receptors in the trimer-of-dimers arrangement (Fig. S1). In addition, the translational motions of receptors in CSUs are probably constrained by their periplasmic domains and by tight interactions between their hairpin tips and the CheA/CheW signaling proteins (46, 54) (Fig. S1). We assume, therefore, that crosslinking in single-CYS receptors reflects dynamic motions within a homodimer rather than between the neighboring receptor molecules of core signaling units.

The MH1 and MH2 helices behaved comparably with respect to *e*-CYS reporters (Fig. 2*A*). In cells not pre-exposed to serine, less than 10% of the reporter subunits formed dimeric crosslinking products, implying little dynamic motion of the MH bundle in the absence of a serine stimulus. However, in cells pre-treated with 10 mM serine, three MH1 reporters and three MH2 reporters crosslinked more substantially, ranging up to 20% yield of 1-1’ or 2-2’ products. Overall, *e*-residue sites at each end of the MH bundle exhibited little or no dynamic behavior either with or without serine pretreatment, whereas *e* sites in the middle of the MH bundle showed serine-enhanced dynamics.

**Fig. 2.**
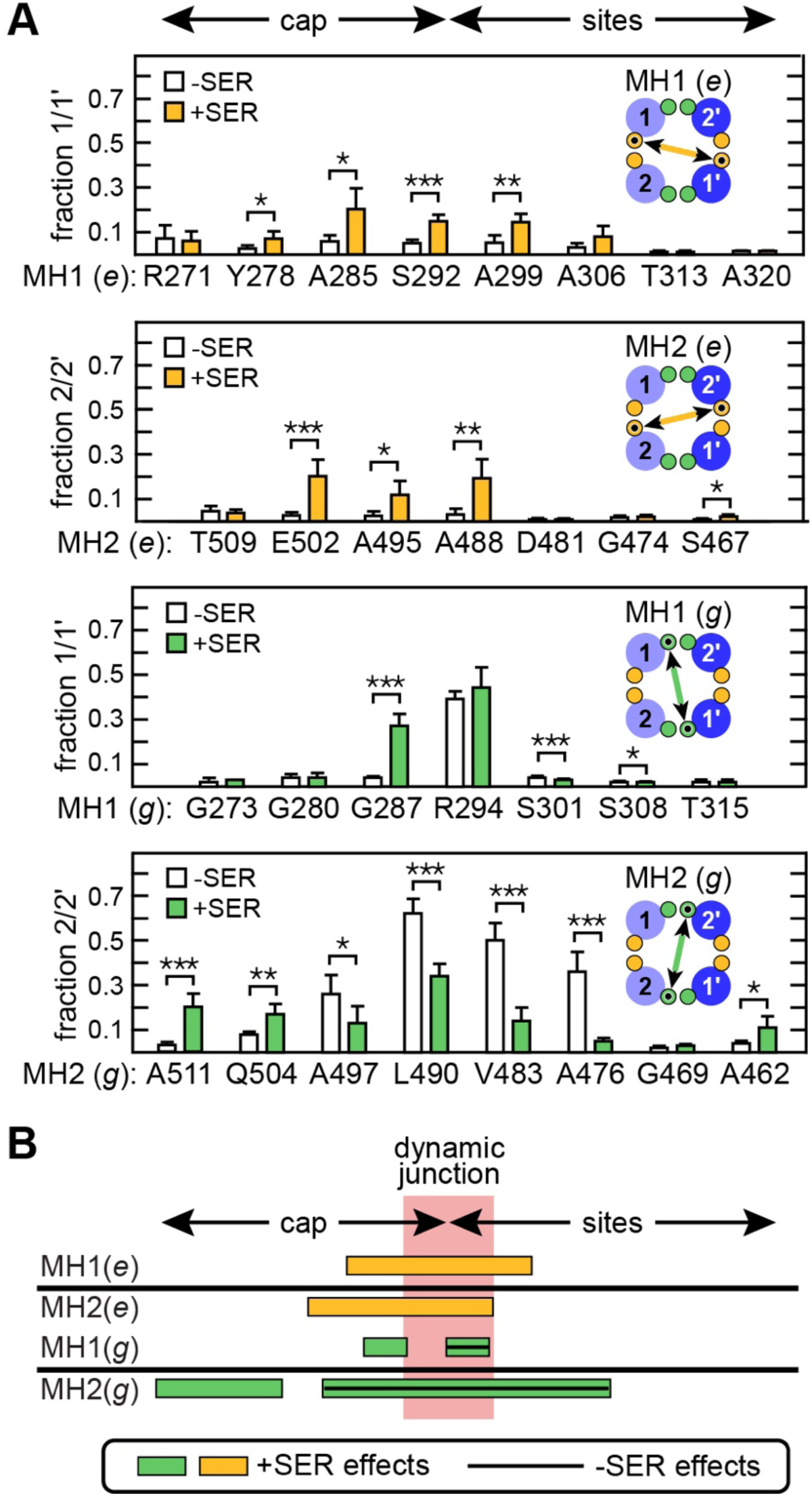
Evidence for a dynamic junction in the Tsr MH bundle. (A) BMOE crosslinking between Tsr subunits with a cysteine reporter at a bundle edge residue. The cartoon insets in each panel depict the subunit helices (light and dark blue), the positions of their *e* (orange) and *g* (green) edge residues and the crosslinked reporter sites (small black circles). Data are means and standard deviations from three or more biological replicates. Asterisks indicate reporter positions with significant ±SER differences (white *vs.* colored bars) as determined by the following *p* values: <0.05 (*), <0.01 (**), or <0.001 (***). (B) Summary of the data in (A). Lines and boxes indicate single-CYS reporters with crosslinking yields of 15% or more in the absence (lines) and/or presence (boxes) of 10 mM serine.

The MH1 *g*-CYS reporter sites at each end of the bundle exhibited low crosslinking with and without serine, much like the MH1 *e*-CYS reporters (Fig. 2*A*). The MH2 *g*-CYS reporters were similarly quiescent at the sites end of the MH bundle, but more dynamic than their MH1 counterparts at the HAMP-proximal cap end of the MH bundle. Both MH1 and MH2 *g*-CYS reporters exhibited substantial crosslinking yields near the cap-sites junction. In MH1, the G287C reporter showed more than a five-fold crosslinking increase in the presence of serine and the adjacent R294C reporter exhibited over 40% crosslinking both in the absence and presence of serine (Fig. 2*A*). Three MH2 *g*-CYS reporters spanning the cap-sites junction (L490C, V483C, A476C) showed 35-60% crosslinking in the absence of serine and pretreatment with serine reduced their crosslinking propensities by two-fold or more (Fig. 2*A*).

These single-CYS crosslinking results indicate that serine stimuli differentially influence the dynamic behaviors of reporters in the MH1 and MH2 helices. The dynamics of both helices were most pronounced at the junction of the MH cap and sites segments, which we designate the dynamic junction of the MH bundle (Fig. 2*B*). At the dynamic junction, serine enhanced crosslinking of the MH1 *g*-CYS reporter (R294C) and both MH1 (S292C) and MH2 (A288C) *e*-CYS reporters but suppressed crosslinking of L490C and other MH2 *g*-CYS reporters spanning the junction. Elevated crosslinking behaviors spread outward from the dynamic junction, mainly toward the HAMP-proximal cap end of the MH helices. The HAMP-distal sites end of the MH bundle exhibited no discernable dynamic behavior in single-CYS reporters. As we describe below, we interpret these results to show the dynamic junction plays a unique mechanistic role in signal transmission through the Tsr MH bundle and may account for the MH bundle structural dynamics previously documented with in vitro studies of the Tar receptor.

### Dynamic Motions Cause BMOE-Dependent Mobility Shifts in Tsr Monomers

During single-CYS crosslinking experiments, we noted that upon BMOE treatment many Tsr reporters produced a new SDS-PAGE band just above or below the monomer position (Fig. S2*A*). Bandshifts occurred for both MH1 and MH2 single-CYS reporters and many of those evinced serine-enhanced effects (Fig. S2*A*). Because the affected reporter sites spanned the dynamic junction and often exhibited serine-enhancement, we suggest that these effects arise from dynamic behaviors that provide BMOE access to the subunit reporter sites (summarized in Fig. S2*B*).

Single amino acid replacements in MCP molecules can also produce modest shifts in subunit mobility in SDS-PAGE experiments, possibly by the same mechanism. The best examples are the effects of adaptational modifications: E residues at adaptation sites slow electrophoretic mobility, whereas methylated E or methyl-mimic Q residues speed mobility (55–58). The mechanism behind these bandshifts is unclear but may reflect differences in the overall density of negatively-charged SDS molecules that coat the receptor subunits (59). Thus, negative charges at key receptor positions might lead to slower gel migration by interfering with SDS binding. BMOE might produce a similar mobility shift by occasionally crosslinking an accessible cysteine in a Tsr subunit to glutathione or some other small, negatively-charged cytoplasmic component that bears a maleimide-reactive sulfhydryl group.

Cu^2+^-phenanthroline (CuPhen) treatment also produced monomer mobility shifts and dimer crosslinking products (Fig. S2*C*). In 100-second reactions CuPhen and BMOE produced similar extents of dimeric crosslinking products, but the shifted monomer products accumulated more slowly under CuPhen conditions, implying that the shift effect involves an intrinsically slower reaction than does subunit-subunit crosslinking (Fig. S2*C*). The band-shifted CuPhen species reverted to monomers upon treating the samples with a reducing agent, but reducing treatment had little effect on the band-shifted monomers produced by BMOE (Fig. S2*C*). These results are consistent with the possibility that disulfide (CuPhen) or maleimide (BMOE) linkage to a thiol-bearing compound like glutathione produces the band-shifted species. We conclude that the monomer bandshift effect provides a fortuitous and reproducible means of assessing receptor dynamic behavior independently of subunit-subunit crosslinking. Both readouts identify similar dynamic regions in the MH1 and MH2 helices that span the dynamic junction (Fig. S2*B*).

### Signaling Behaviors of Single-CYS Tsr Reporters

We used an in vivo assay based on Förster resonance energy transfer (FRET) (60–62) to assess the CheA-control signaling behaviors of the Tsr single-CYS receptors. This assay measures interaction between fluorophore-tagged CheY (yellow fluorescent protein, YFP) and its phosphatase partner CheZ (cyan fluorescent protein, CFP). CheA-mediated phosphorylation of CheY-YFP promotes binding with CheZ-CFP, producing a FRET signal that reflects cellular phospho-CheY levels and, in turn, the receptor-controlled kinase activity of CheA. Serine dose-response experiments were conducted in host strain UU2567 (62), a close relative of the UU2610 strain used for the single-CYS crosslinking surveys. In that strain, which lacks the sensory adaptation enzymes, Tsr in the wild-type [QEQEE] modification state inhibits 50% of CheA activity at 15-20 µM serine, its *K1/2* response value (16, 21, 22, 24, 62–65).

Tsr *e*-CYS reporters were generally OFF-shifted in FRET assays, with only one receptor showing a serine response threshold above the wild-type (Fig. S2*D*). In contrast, *g*-CYS receptors were generally ON-shifted in FRET assays, with all but two above the wild-type *K_1/2_* (Fig. S2*D*). These results show that cysteine replacements at *g*-position residues shift Tsr output toward the ON state and replacements at *e*-position residues shift output toward the OFF state. These mutant behaviors imply that the native *g* residues of the Tsr MH bundle play important roles in promoting kinase-OFF output, whereas *e* residues are important for kinase-ON output.

### Crosslinking Surveys of CYS-Pair Receptors in the Tsr MH Bundle

To investigate how serine-induced conformational signals propagate through the Tsr dynamic junction, we followed in vivo BMOE-promoted crosslinking between pairs of cysteine replacements at heptad edge-residue (*e*, *g*) positions. Our working model posits that serine stimuli induce axial helix rotations that shift the MH bundle from a conventional “knobs-in-holes” *a-d* packing arrangement in the ON state to a complementary *x-da* OFF-state arrangement in which the edge residues at *g* positions rotate toward the bundle core and edge residues at *e* positions rotate away from the packing core (Fig. 3*A*) (16). We constructed Tsr reporters bearing adjacent pairs of *g*-CYS sites (designated *g/g* or *g/g*-CYS receptors; Table S1; Fig. S3*B*). Note that adjacent *g*-position residues in the MH1 and MH2’ helices have a staggered arrangement (Fig. S3). Such *g/g* reporter sites would be expected to move closer and crosslink more readily in the presence of serine, according to our working model (Fig. 3*A*).

**Fig. 3.**
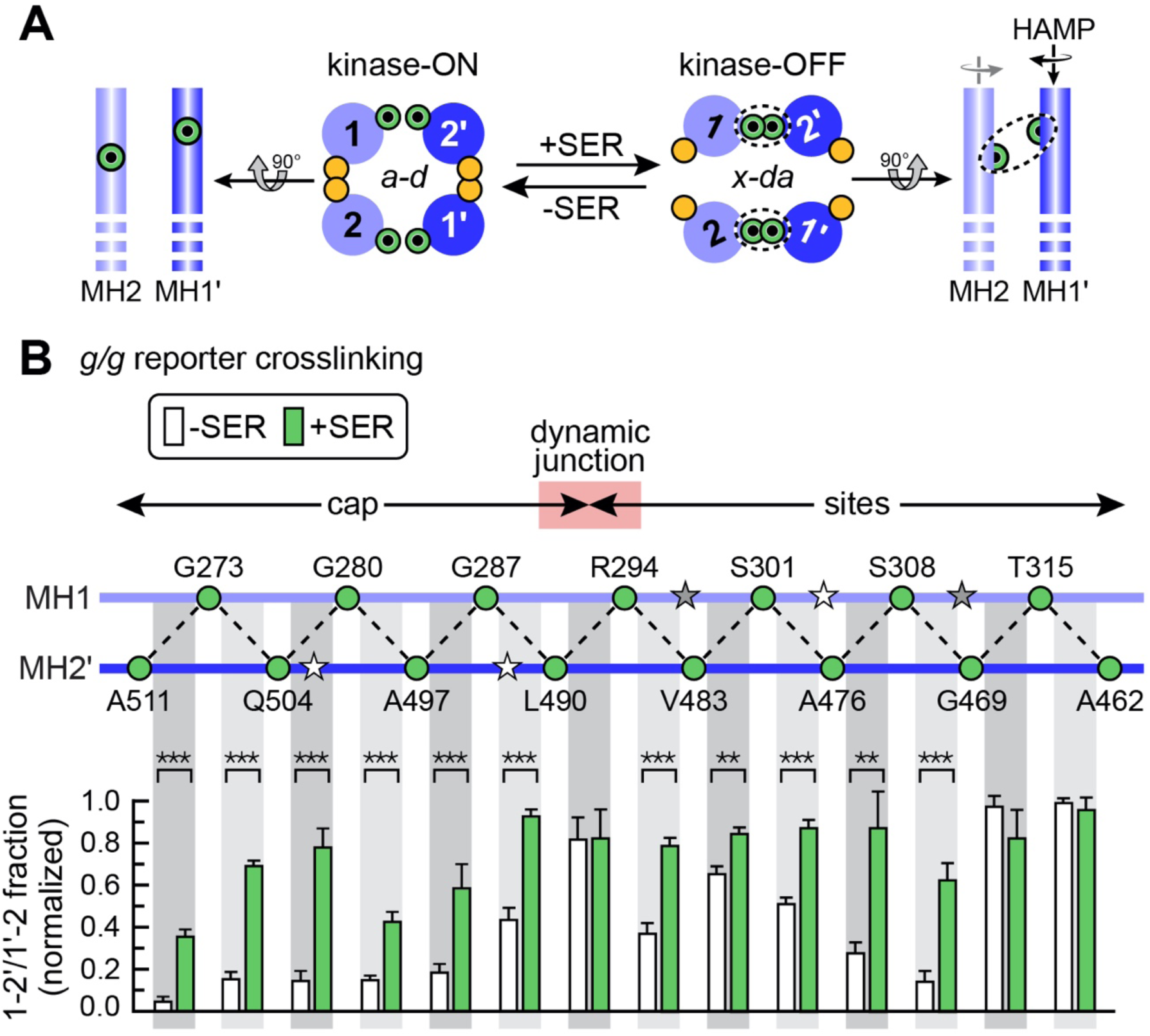
BMOE crosslinking scan of *g/g* reporter pairs in the Tsr MH bundle. (A) Working model of MH bundle packing and signaling configurations. The cartoons depict the subunit helices (light and dark blue), the positions of their *e* (orange) and *g* (green) edge residues and the cysteine reporter sites (small black circles). Serine-induced axial rotation is predicted to enhance crosslinking of *g/g* reporter pairs by shifting the MH bundle from *a-d* (ON) to *x-da* (OFF) packing (16). (B) *g/g* cysteine pairs surveyed. Cysteine replacements were created at each of the indicated *g-*position residues, then combined to make Tsr subunits bearing adjacent MH1/MH2’ *g/g* pairs (dashed lines). Stars represent wild-type Tsr adaptation site residues (white, E; gray, Q) that were not manipulated in these experiments. To simplify the data presentation, sites are shown evenly spaced between the helices, but darker gray shading indicates the longer distance *g/g* pairs. See Fig. S4 for a more accurate structural view. Crosslinking data are means and standard deviations for doubly-crosslinked (1-2’/1’-2) products from three or more biological replicates normalized to the highest value (T315C/A462C -SER). Raw data values for these experiments are given in Table S2. Asterisks indicate reporter positions with significant ±SER differences as determined by the following *p* values: <0.01 (**) or <0.001 (***).

We tested *g/g* crosslinking behaviors with the same BMOE reaction conditions used for the single-CYS receptors. (See Fig. S4 *A* and *B* for representative SDS-PAGE band patterns and quantitative analyses of the crosslinked products. Note that there are no band-shifted monomers in the gels, confirming that the dimeric crosslinking reactions are much faster.) Under these conditions serine produced highly significant increases in the 1-2’/1’-2 doubly crosslinked BMOE product for all six MH cap reporter pairs and for five reporter pairs in the MH sites region (Fig. 3 *A* and *B*). Three reporter pairs (R294C/L490C at the dynamic junction and T315C/G469C and T315C/A462C at the HAMP-distal end of the MH sites segment) exhibited comparably high crosslinking signals in the absence, as well as presence, of serine (Fig. 3*B*). Crosslinking timecourses of these highly reactive *g/g* reporter pairs at a 10-fold lower BMOE concentration showed that their doubly crosslinked product formed more rapidly in serine-exposed cells than it did in buffer-treated controls (Fig. S5*C-E*). Thus, *g/g* reporters throughout the MH bundle underwent demonstrable signaling-related structural changes consistent with serine-induced *a-d* to *x-da* packing transitions at the reporter sites.

A key prediction of this signaling model not previously tested is that adjacent *e-*position cysteines in MH1 and MH2, which also have a staggered arrangement (Fig. S3), should crosslink more readily in the absence of serine because *e/e* reporter sites lie closer in the *a-d* bundle configuration than in the serine-promoted *x-da* packing arrangement (Fig. 4*A*). Accordingly, we surveyed a corresponding set of *e/e* reporter pairs across the Tsr MH bundle (Fig. 4). These should (Fig. 4*A*) and did (Fig. S5) form monomer-sized crosslinking products. Initial experiments done with BMOE reaction conditions optimal for *g/g* reporters produced high levels of *e/e* crosslinking in both the absence and presence of serine. Based on reaction timecourses at lower BMOE concentrations (Fig. S6 *A* and *B*), we chose a 10-fold lower BMOE concentration (20 µM), a 10-fold shorter reaction time (10”), and a lower acrylamide concentration (8%) to better resolve the *e/e* crosslinking products and serine-related effects. Some *e/e*-crosslinked subunits migrated slower in SDS-PAGE immunoblots than did uncrosslinked Tsr subunits; others migrated faster, depending on the positions of the reporter sites along the MH bundle (Fig. S5*C*).

**Fig. 4.**
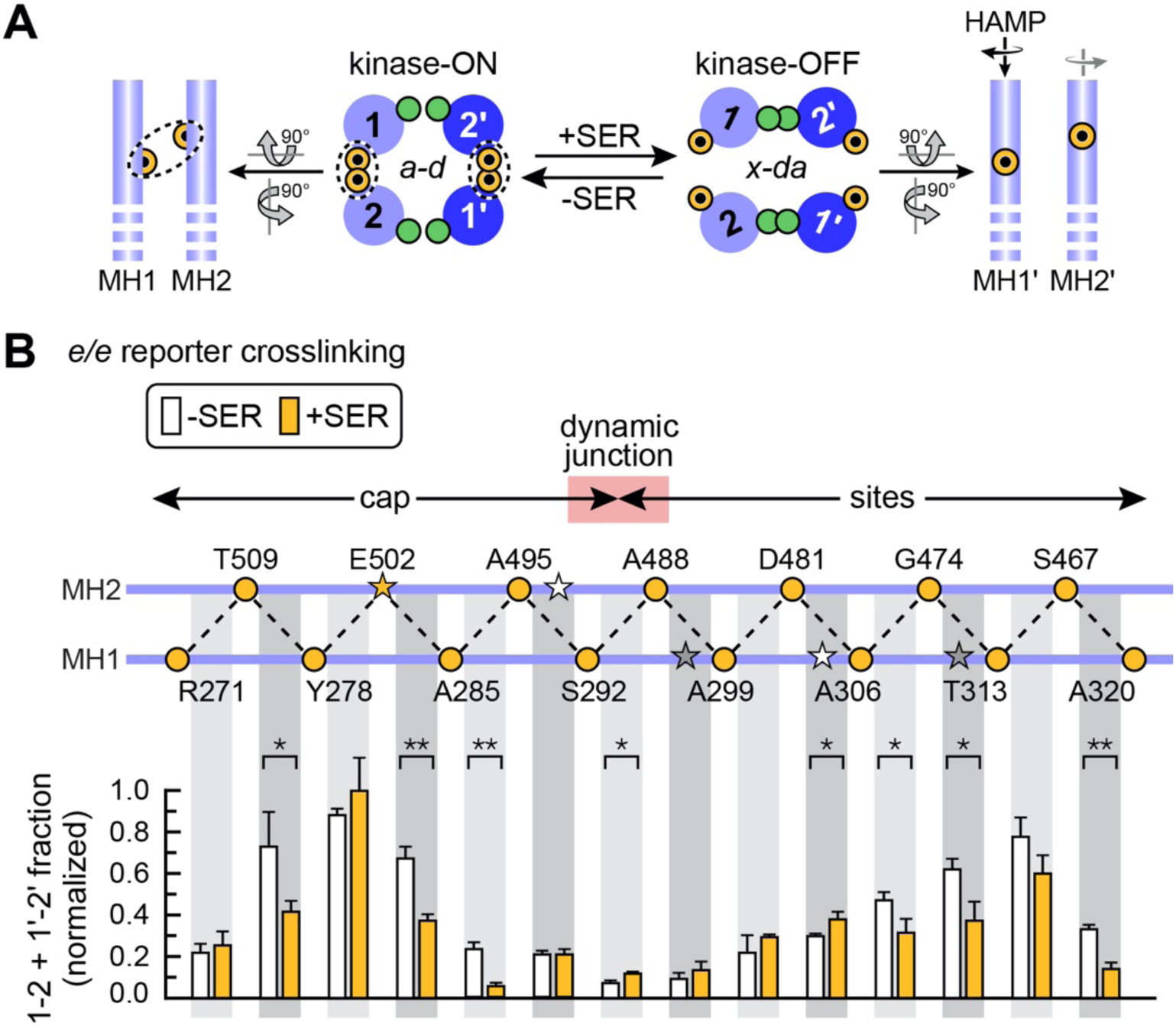
BMOE crosslinking scan of *e/e* reporter pairs in the Tsr MH bundle. (A) Working model of MH bundle packing and signaling configurations. The cartoons depict the subunit helices (light and dark blue), the positions of their *e* (orange) and *g* (green) edge residues and the crosslinked reporter sites (small black circles). Serine-induced axial rotation of the MH1 helices is predicted to inhibit crosslinking of *e/e* reporter pairs by shifting MH bundle packing toward an *x-da* (kinase-OFF) configuration, whereas adjacent *e/e* reporter sites should be closer and more readily crosslinked in an *a-d* (kinase-ON) packing arrangement. (B) e*/e* cysteine pairs surveyed. Cysteine replacements were created at each of the indicated *e-*position residues, then combined to make Tsr subunits bearing adjacent *e/e* pairs (dashed lines). Stars represent wild-type Tsr adaptation site residues (white, E; gray, Q) that were not manipulated in these experiments. To simplify the data presentation, sites are shown evenly spaced between the helices, but darker gray shading indicates the longer distance *e/e* pairs. See Fig. S4 for a more accurate structural view. Crosslinking data are means and standard deviations for intra-subunit crosslinking products (1-2 and 1’-2’) from three or more biological replicates normalized to the highest value (Y278C/E502C +SER). Raw data values from these experiments are given in Table S2. Asterisks indicate reporter positions with significant ±SER differences as determined by the following *p* values: <0.05 (*) or <0.01 (**).

The *e/e* reporter pairs showed substantial differences in the extent of crosslinking through the MH bundle (Fig. 4*B*). Six reporter pairs (three in the cap and three in the sites region) crosslinked significantly more in the absence than in the presence of serine (Fig. 4*B*), confirming that the serine-induced bundle arrangement slows crosslinking between CYS reporter sites at those MH1/MH2 *e*-heptad positions. Reaction timecourses might reveal significant serine-dependent crosslinking differences for other *e/e* CYS pairs, but we did not pursue that issue in the present study.

### BMOE Crosslinking Shifts Tsr *g/g* Receptors toward Kinase-OFF Output and *e/e* Receptors toward Kinase-ON Output

In FRET kinase assays, the behaviors of CYS-pair reporters were like, but in some cases more extreme than, their single-CYS components (Table S1). Six *g/g* reporters exhibited an elevated *K1/2* value, consistent with ON-shifted signaling properties. Eight other *g/g* receptors failed to respond to even 10 mM serine, but their FRET signal dropped substantially when the cells were challenged with 3 mM potassium cyanide (KCN). KCN treatment collapses cellular ATP levels, the phosphodonor for CheA autophosphorylation, and consequently prevents CheA-mediated phosphorylation of CheY (62) (Fig. S6*A*). By contrast, eight *e/e* reporters were kinase-OFF and five others had OFF-shifted serine responses with *K1/2* values below the wild-type (Table S1). As with the single-CYS reporters, these mutant output behaviors imply that the native Tsr *g* and *e* residues of the MH bundle play important roles in promoting kinase-OFF and kinase-ON CheA outputs, respectively.

The MH1 helix-rotation model predicts that because BMOE crosslinks are irreversible, crosslinking should trap the helices of Tsr *g/g* receptors in the rotated state and reduce or eliminate their kinase activity. This effect proved true for 4/5 MH cap *g/g* receptors in our earlier study (16). That trend prevailed in 8/9 additional MH bundle *g/g* reporters, which were driven to lower *K1/2* or fully kinase-OFF behavior by BMOE exposure in the FRET assay (Fig. S6). The one exception was the S308C/G469C reporter near the HAMP-distal end of the MH bundle. This receptor remained locked in kinase-ON output after challenge with a combination of serine plus BMOE (Fig. S6*A*), similar to one serine non-responsive *g/g* reporter in the cap (16).

We examined five *e/e* reporters for BMOE signaling effects in the FRET assay; all five showed behaviors consistent with the prediction that *e/e* crosslinks would shift output toward the kinase-ON state (Fig. S7). BMOE abolished serine responses of the Y278C/E502C receptor and locked it in kinase-ON output (Fig. S7*A*). Remarkably, two kinase-OFF reporters (R271C/T509C, S292C/A495C) became kinase-active following BMOE exposure and readily responded to small serine stimuli (*K1/2* values below 1 µM) (Fig. S7*B*). Two other kinase-active, serine-responsive receptors (Y278C/T509C, A306C/G474C) retained kinase activity after BMOE treatment but became much less sensitive to serine (>100-fold higher *K1/2* values), again consistent with a crosslinking-induced shift toward the ON output state (Fig. S7*C*).

These results demonstrate a relationship between the extent of BMOE crosslinking and the degree to which BMOE treatment shifted *g/g* receptors toward kinase-OFF output and *e/e* receptors toward kinase-ON output. The Tsr molecules in our crosslinking and FRET experiments assemble arrays of cooperatively networked signaling teams, so not all cell receptors would necessarily need to be crosslinked to lock array output in an ON or OFF state. Conversely, owing to variable sizes and connectivities of receptor signaling teams within arrays, even extensively crosslinked arrays could retain some kinase activity after BMOE treatment.

### Crosslinking and Kinase-control Behaviors of CYS-Pair Receptors

A comparison of the crosslinking and kinase-control behaviors of CYS-pair reporters shows that these two signaling readouts differ in stimulus sensitivity (Fig. 5). Seven *e/e* reporters spanning the dynamic junction were locked-OFF in FRET assays, indicating that their hairpin tip produced no detectable CheA activity. Yet reporters at each end of the dynamic junction exhibited significant serine-dependent crosslinking effects: serine inhibited crosslinking of the E502C/A285C and A285C/A495C cap reporters and enhanced crosslinking of the D481C/A306C sites reporter (Fig. 4). Evidently, the locked-OFF conformation at the hairpin tip does not reflect the reporter conformation in the MH bundle, which is still responsive to a serine stimulus. The disparity between crosslinking and FRET readouts was even more striking for the *g/g* reporters: eight exhibited serine non-responsive, locked-ON kinase activity in FRET assays, yet serine treatment substantially increased crosslinking for six of them, from Q504C/G280C in the cap to A476C/S308C in the sites (Fig. 3; Fig. 5). These findings demonstrate that in vivo crosslinking assays can provide a more sensitive structural readout of stimulus response changes than does the FRET kinase assay.

**Fig. 5.**
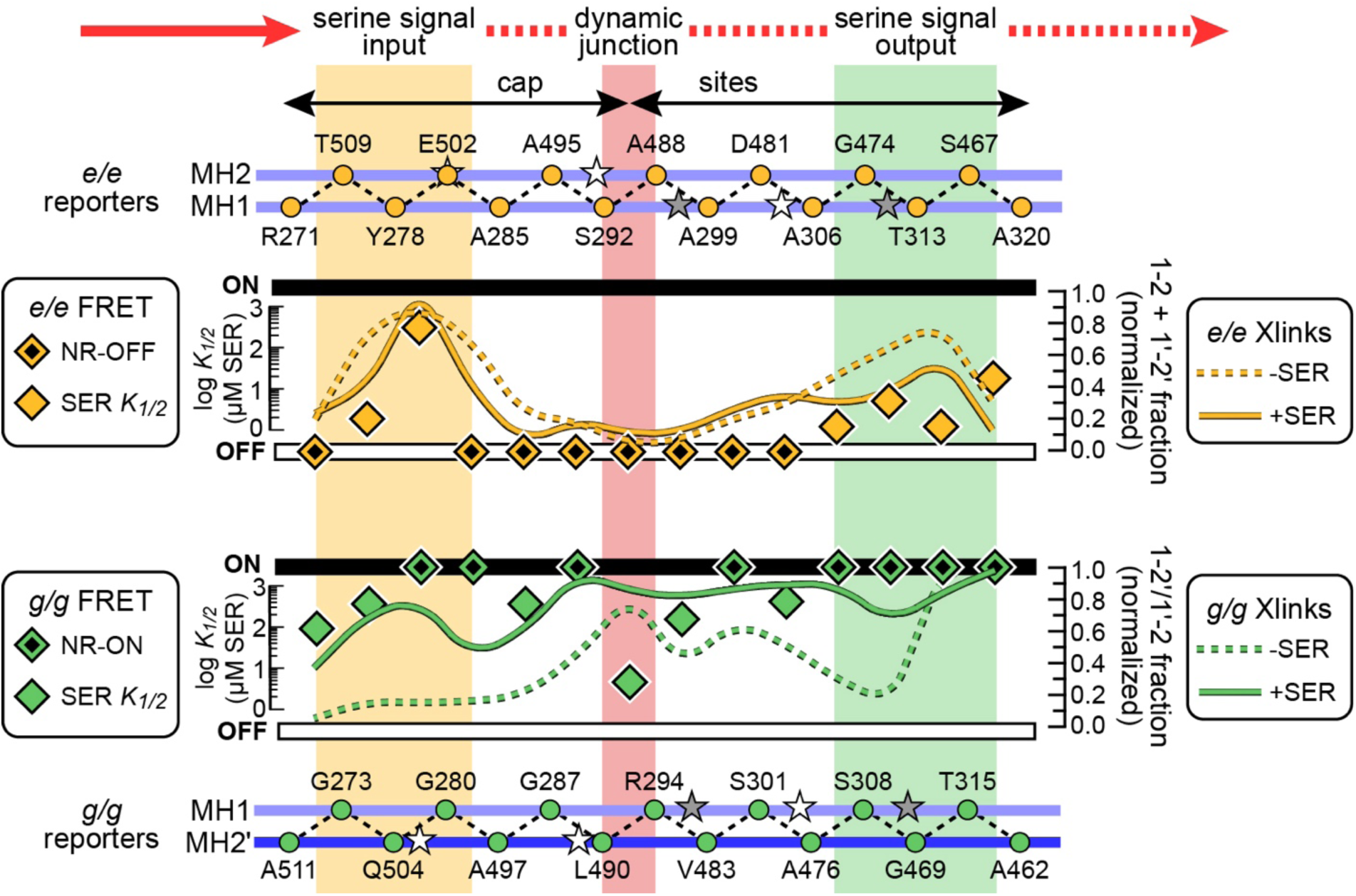
Comparison of crosslinking and kinase-control behavioral readouts for CYS-pair reporters. These data illustrate behavioral differences between the serine signal input and output segments of the MH bundle and between crosslinking and FRET kinase behaviors throughout the bundle. The helix cartoons at top and bottom depict the subunit helices (light and dark blue) and the positions of their *e* (orange) or *g* (green) residues. Stars indicate the approximate positions of the wild-type E (white) and Q (gray) adaptation sites that were not manipulated in these experiments. Normalized crosslinking values (UU2610 host; CheRB^-^) are summarized by smoothed lines for *e/e* CYS-pair receptors from Fig. 4B (upper panel) and *g/g* CYS-pair receptors from Fig. 3B (lower panel). Diamonds show FRET values for <u>uncrosslinked</u> receptors from Table S1 (UU2567 host; CheRB^-^). NR-OFF receptors exhibited no response to serine stimuli up to 10 mM and no FRET change in response to 3 mM KCN, indicative of no CheA activity (62). NR-ON receptors exhibited kinase activity upon KCN challenge (62), but failed to inhibit that activity when challenged with 10 mM serine.

Our crosslinking assays effectively produce short-exposure snapshots of MH bundle packing arrangements. These conformational readouts detect helix-packing responses to attractant stimuli within individual receptor dimers and do not depend on the higher-order interactions of receptors in core signaling units or arrays (16, 66). In a two-conformation (*a-d* versus *x-da*) model of Tsr MH bundle output states the extent of *e/e* crosslinking in the presence of serine should reflect a receptor’s ON conformational bias at the reporter sites. Conversely, the extent of *g/g* crosslinking in the absence of serine should reflect a receptor’s OFF conformational bias at the reporter sites. Although the *e/e* and *g/g* data were obtained with different reaction conditions, their normalized values permit useful comparisons between them.

The conformational bias estimates revealed three MH bundle regions with distinctive signaling behaviors: The HAMP-proximal (serine signal input) end of the bundle had ON-biased conformation; the HAMP-distal (serine signal output) end of the bundle had moderate OFF-biased conformation. The dynamic junction showed a strongly OFF conformational bias: high *g/g* crosslinking levels in the presence and absence of serine and very little *e/e* under either condition. The high OFF bias at the dynamic junction that progressively diminished through the sites segment until again peaking at the signal output end of the MH bundle (Fig. 5).

### AlphaFold Insights into MH Bundle Signaling Structures

We generated atomic structures of the full-length Tsr dimer with AlphaFold 3 (67) and analyzed their MH bundle packing arrangements with SamCC Turbo, which computes Crick angles (sidechain orientations) in 4-helix coiled-coil bundles (68). The MH2 helices showed Crick angle deviations characteristic of *x-da* packing over residues 486-494 spanning the dynamic junction (Fig. 6*A*). The corresponding segment of the MH1 helices showed a similar but less pronounced rotation trend (Fig. 6*A*). Rotation of the MH2 helix into an *x-da* packing orientation at the dynamic junction coincided with a pronounced bulge and kink (Fig. 6*B*), similar to a feature noted in the crystal structure of Tm14, a soluble MCP of *Thermotoga maritima* (69). The Crick angle deviations indicate the input and output ends of the MH bundle had *a-d* packing configurations, whereas the dynamic junction showed *x-da* packing (Fig. 6 *C* and *D*). These structural features suggest the dynamic junction adopts an *x-da* packing arrangement in the absence of a serine stimulus and that rotation of the MH2 helix is probably the principal driving force for that configuration, which would explain the very high crosslinking of MH2 *g*-CYS reporters that span the dynamic junction. These findings provide a structural context for the unique crosslinking behaviors of the dynamic junction and suggest a mechanism for signal propagation through the MH bundle, as discussed below.

**Fig. 6.**
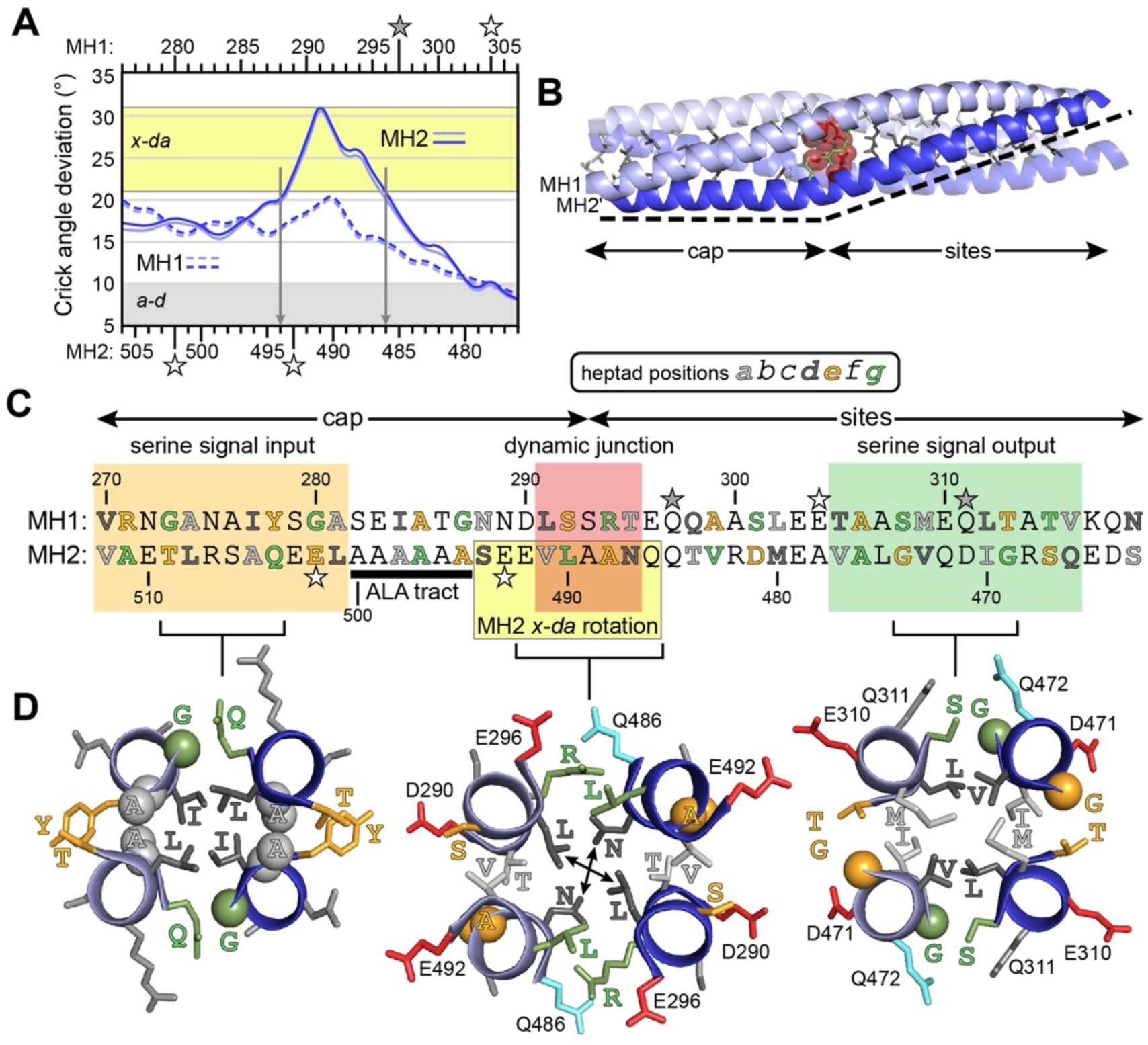
Signaling-related AlphaFold 3 structures of the MH bundle and dynamic junction. (A) Crick angle deviations (axial rotations) of MH1 and MH2 residues from a SamCC Turbo analysis (68) of atomic coordinates for a full-length Tsr dimer model generated by AlphaFold 3 (67). The *a-d* packing configuration is defined by Crick angle deviations ±10° (gray), whereas *x-da* packing (yellow) is shown here as an axial rotation of 26 ± 5° (89). (B) Structure of the MH bundle in an AlphaFold 3 model of the Tsr dimer. Side-chains of residues at *a* (light gray) and *d* (dark gray) positions are shown as sticks. Semi-transparent spheres (red) represent side-chain atoms of the dynamic junction *d*-heptad residues L291 and N487. Green sticks show side-chains of dynamic junction *g*-heptad residues R294 and L490. The dashed line traces a pronounced bulge and kink at the dynamic junction. (C) Primary structure and key regions of the Tsr MH bundle. (D) One-heptad slab views of packing configurations in the AlphaFold 3 Tsr model. Key residues are labeled and color-coded to the sequence in (C) and negatively charged residues are shown in red. Glycine residues are shown as a sphere; Cα and Cβ atoms of alanine sidechains otherwise difficult to discern are also shown as spheres. Arrows in the dynamic junction structure indicate potential interactions between the *x* positions in the *x-da* packing configuration. Methylation site Q311, and perhaps the other cyan-colored glutamine residues, influence MH bundle packing arrangements and stabilities.

## DISCUSSION

### Signaling-related conformations in the Tsr MH bundle

Three lines of evidence indicate that *x-da*-like packing arrangements of the Tsr MH bundle produce kinase-OFF output: (i) Residues at adjacent MH1 and MH2 *g*-heptad positions lie closer in the *x-da* arrangement than they do in the *a-d* configuration (Fig. 3*A*) and OFF-shifting serine stimuli enhanced BMOE crosslinking of *g/g* reporter pairs (Fig. 3*B*). (ii) Most *g*-position cysteine replacements shifted Tsr output toward the ON state (Fig. S2*D*; Fig. 5; Table S1), suggesting that destabilization of the packing arrangement promoted by native *g*-position residues reduces kinase-OFF output. (iii) BMOE crosslinking of *g/g* reporter pairs shifted their output back toward the kinase-OFF state (Fig. S6 *B-D*).

Analogous evidence indicates that *a-d*-like packing configurations of the Tsr bundle produce kinase-ON output: (i) Residues at adjacent MH1 and MH2 *e*-heptad positions lie closer in the *a-d* arrangement than they do in the *x-da* configuration (Fig. 4*A*) and OFF-shifting serine stimuli slowed BMOE crosslinking of *e/e* reporter pairs (Fig. 4*B*). (ii) Most *e*-position cysteine replacements shifted Tsr output toward the OFF state (Fig. S2*D*; Fig. 5; Table S1), suggesting that destabilization of the packing arrangement promoted by native *e*-position residues reduces kinase-ON output. (iii) BMOE crosslinking of *e/e* reporter pairs shifted their output back toward the kinase-ON state (Fig. S7).

### A Three-state Signaling Model for the Tsr MH Bundle

The crosslinking behaviors of CYS-pair receptors indicate that signaling through the Tsr MH bundle can be understood in terms of a three-state structural model involving kinase-ON (*a-d*) and kinase-OFF (*x-da*) helix-packing arrangements linked through unstable or unstructured intermediates (Fig. 7). The HAMP domain, which provides conformational input to the MH bundle, probably has analogous signaling modes (23, 70–72), but we do not consider those here. A four-residue phase stutter couples the HAMP AS2 helices to the MH1 helices in opposed packing arrangements, such that an *x-da* packed HAMP bundle should favor *a-d* packing of the MH cap, whereas an *a-d* HAMP bundle should favor *x-da* packing of the MH cap (16, 24, 72, 73) (Fig. 7). Thus, interconversion of the *a-d* and *x-da* configurations in the signal input and output ends of the MH bundle occurs through HAMP-promoted axial rotation of the MH1 helices and subsequent counter-rotation of the MH2 helices to optimize stability at their packing interfaces (16, 73).

**Fig. 7.**
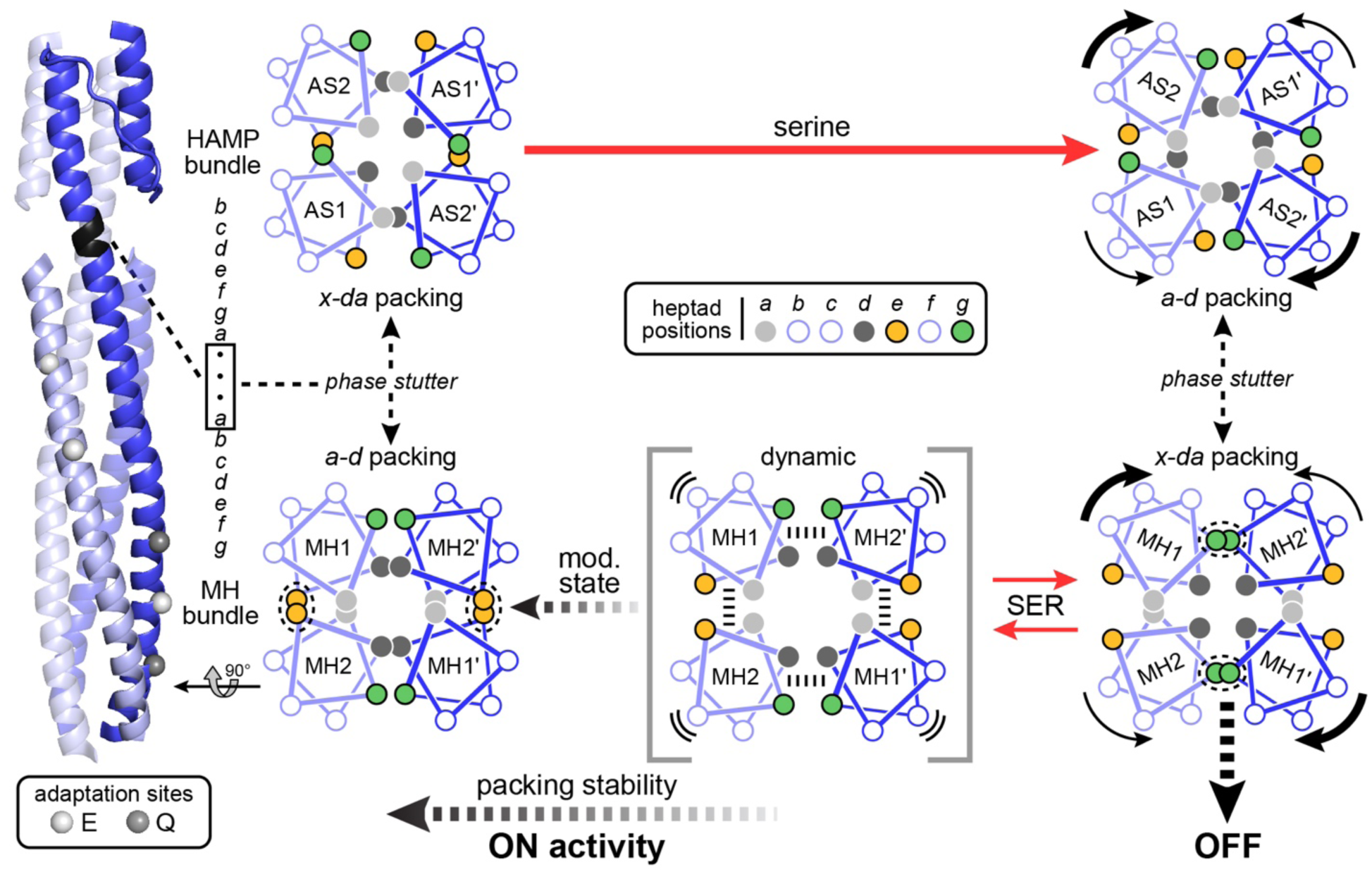
Three state model of MH bundle signaling in Tsr. The ribbon diagram at left shows subunits of the HAMP and MH bundles of a Tsr dimer extracted from a model of the core receptor signaling unit (54) with helices at the rear dimmed and colors keyed to the helical wheels to the right. Spheres depict the alpha carbons of E (white) and Q (gray) adaptation site residues. The 4-residue phase stutter (black) joins the AS2 and MH1 helices ∼26° out of register. For simplicity, the first three residues of the stutter helix lack a heptad designation (•); the last stutter residue (V267) occupies an *e*-heptad position in the AS2 register, but an *a*-heptad position in the MH1 register. Helical wheel diagrams depict top-down cross-section views through the bundles. Due to the phase stutter, an *a-d* packed HAMP bundle favors an *x-da* packed MH bundle, whereas an *a-d* packed MH bundle favors *x-da* packing of HAMP. Black curved arrows indicate the directions of helix rotations; thick curved arrows indicate the serine-imposed helix rotations through the phase stutter connections. Dashed lines between helices in the dynamic intermediate state represent the principal helix-helix interactions that modulate MH bundle packing stability. In the presence of serine (SER) the MH bundle is in rapid equilibrium between *x-da* packing and dynamic states. The adaptational modification state of Tsr (mod. state) spans a range from fully unmodified (5E) to fully modified (5Q and/or methylated E) residues per subunit.

In addition to conserved, but often atypical, *a* and *d* packing residues throughout the MH bundle that could moderate helix pairing interactions, conserved acidic residues on solvent-exposed faces of the MH1 and MH2 helices (Fig. S8) also contribute to MH bundle dynamics. These acidic faces create an extended negatively-charged surface that could contribute to bundle dynamics by reducing helix stability (33). Three of the MH1 acidic-face residues are sensory adaptation sites, where methylation or a neutral sidechain replacement would reduce overall negative charge on the acidic face and thereby enhance MH1 helicity and packing interactions. Indeed, in vitro studies of full-length Tar dimers in nanodiscs and soluble Tar signaling fragments found that MH1 was more dynamic and less helical in the [EEEE] state than in the [QQQQ] modification state (14, 38). The MH2 helix also has solvent-exposed acidic faces flanking the dynamic junction that might modulate helix stability in a similar fashion (33) (Fig. S8). Two of the residues in the acidic face that spans the MH2 alanine tract (24) are sites of adaptational modifications in Tsr (63, 74).

Our three-state signaling model proposes that the *a-d* and *x-da* packing states are in equilibrium with more dynamic bundle configurations, with transition rates controlled by both chemoeffectors and by adaptational modifications (Fig. 7). We suggest that adaptational modifications mainly impact the helix stability needed for packing interactions common to both the *a-d* and *x-da* bundle arrangements, whereas serine-induced helix rotations specifically alter the relative strengths of intra- and inter-subunit helix interactions, thereby promoting a bundle-packing geometry (*x-da*) that is seldom accessed through dynamic motions alone at any modification state, with the notable exception of the dynamic junction.

### Signaling Role of the Dynamic Junction

Tar receptors with single-CYS reporters form crosslinks in vitro over long-duration CuPhen treatments at essentially any MH bundle residue position (29, 34). By contrast, in vivo BMOE crosslinking assays with Tsr reporters capture short-exposure snapshots of the predominant dynamic motions in the MH bundle helices, more clearly reflecting the relative stabilities of their various pairing interactions. Single- and double-CYS reporters crosslinked most readily at a “dynamic junction” between the MH cap and sites segments (Fig. 2; Fig. 5). These behaviors are consistent with the MH1 helix dynamics documented in Tar (14, 37, 38). In addition, Bartelli and Hazelbauer noted that the highly mobile component at one of their Tar spin-label positions was three to four-fold greater than at either flanking reporter site (37). That highly dynamic reporter site (Tar-284/Tsr-286) abuts the Tsr dynamic junction and rotated MH2 segment (Fig. 6*C*), implying that Tar also has a dynamic junction at its cap-sites border.

Several lines of evidence indicate that the Tsr dynamic junction favors an *x-da* packing arrangement in the absence of a serine stimulus. First, the unliganded AlphaFold 3-generated Tsr structure exhibited MH2 rotation and *x-da* packing at the dynamic junction (Fig. 6). The conserved MH2 alanine tract residues (Fig. 6; Fig. S8) might facilitate MH2 Crick angle transitions from *a-d* packing in the MH cap to *x-d*a packing at the dynamic junction. The conserved LLF motif cap residues (Fig. S8) might distort the geometry of *a-d* packing layers in the cap (24) to facilitate the transition to *x-da* orientation through the alanine tract. Second, single-CYS and CYS-pair reporters at the dynamic junction *g*-position residues (R294, L490) crosslinked extensively (Fig. 2; Fig. 3), whereas the corresponding *e*-site reporters (S292, A488) exhibited very little crosslinking in the absence of serine (Fig. 2; Fig. 4). Moreover, the R294 and L490 side-chains are distinctly different from those at *g* positions elsewhere in the MH bundle (Fig. S8). Third, most residues at the dynamic junction are highly conserved and four of them (R294, T295, L490, V491) are unique to Tsr, Tar, and other 36H class MCPs (Fig. S8) (25). Another highly conserved dynamic junction residue, N487 (Fig. 6*D*; Fig. S8), corresponds to the site of the bulge and kink in the Tm14 structure (69).

### Signal Transmission Through the Tsr MH Bundle

At the wild-type [QEQEE] modification state used for all crosslinking assays in the present study, the Tsr dynamic junction seems to be uniquely poised to facilitate serine-induced transitions to OFF-state (*x-da*) packing interactions throughout the MH bundle. In the absence of a serine stimulus, the upstream signal input cap had a strong *a-d* packing bias, whereas the downstream signal output end of the MH bundle had more balanced *a-d* and *x-da* packing modes in evidently rapid equilibrium (Fig. 8). The unstable character of the *x-da* configuration at the intervening dynamic junction extended a bit in both directions through dynamic motions of the MH2 and, to a lesser extent, the MH1 helices (Fig. S2*B*).

**Fig. 8.**
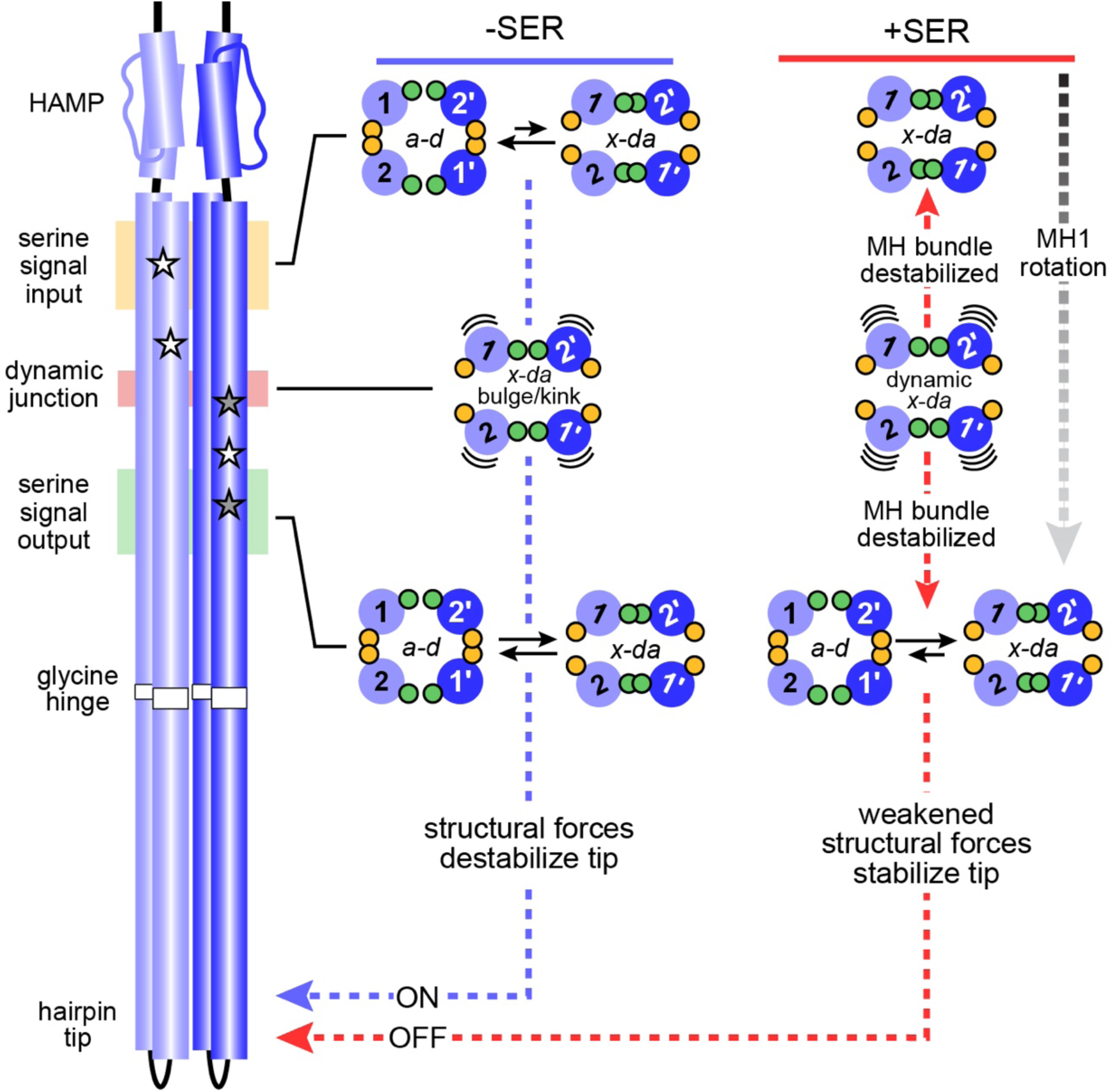
Signal transmission model for the Tsr MH bundle. Cylinders represent alpha-helical segments of Tsr subunits (light and dark blue) of the cytoplasmic domain drawn approximately to scale. Stars indicate adaptational modification site residues, E (white) and Q (gray). Proposed helix-packing configurations in the absence (-SER) and presence (+SER) of a serine stimulus are shown for the signal input, dynamic junction, and signal output regions of the MH bundle. Light and dark blue circles depict simplified top-down cross-section views of the alpha helices with only *e* (orange) and *g* (green) heptad positions shown. The relative lengths of the black arrows indicate the equilibrium bias between *a-d* and *x-da* packing configurations. Curved black lines represent increased dynamic behavior of the helices. MH1 and MH2 residues of the dynamic junction promote MH2 rotation and *x-da* packing at the dynamic junction, which spreads to the input and output segments upon serine-promoted rotation of the MH1 helix. Methylation of the MH1 and MH2 adaptation sites (not shown in this diagram) produces sensory adaptation by enhancing *a-d* packing of the MH bundle (see Fig. 7). Refer to the text for further explanation and discussion of the model.

Serine stimuli produced a dramatic shift toward *x-da* packing throughout the bundle (Fig. 8). The cap adopted an *x-da* configuration with little evidence of *a-d* packing; the bundle output end became *x-da* biased, although it still underwent frequent transitions to *a-d* packing. These changes appear to stem from HAMP-induced axial rotations of the MH1 helices, which probably reinforce MH2 rotation at the dynamic junction and propagate that MH2 *x-da* orientation to the input and output regions of the bundle (Fig. 8). Importantly, the expanded scope of *x-d*a packing promoted by serine does not stabilize overall bundle structure, but rather makes it more dynamic, probably due to the sidechain characters of the MH bundle packing and edge residues (Fig. S8). Equally important, *x-da* packing competes with *a-d* packing, thereby shifting overall bundle output toward the kinase-OFF state. In the presence of serine, packing evidently becomes less stable at the dynamic junction, perhaps reflecting the overall rise in bundle dynamic motions (Fig. 8). This change would account for the enhancing effects of serine on crosslinking of single-CYS reporters at the dynamic junction and its flanking regions (Fig. 2). We suggest that it also accounts for the serine-induced <u>reduction</u> in crosslinking of single-CYS MH2-*g* reporters across the dynamic junction (Fig. 2; Fig. S2 *B*)

### Signal Transmission from the MH Bundle to the Tsr Hairpin Tip

This study further substantiates the dynamic-bundle model of HAMP domain input-output signaling in chemoreceptors, which proposed that attractant stimuli stabilize HAMP structure and destabilize MH bundle structure to transmit kinase-OFF control signals to the receptor tip (19, 23). Our crosslinking results indicate that to reach an effective OFF output state, the MH bundle shifts toward more dynamic *x-da*-packing configurations. The “yin-yang” signaling hypothesis (28) proposed that the MH bundle and hairpin tip are also coupled in structural opposition, in which case an unstable MH bundle should promote a more static hairpin tip.

The conformational and dynamic changes detected in our crosslinking assays occur within individual receptor molecules, and because some CYS-pair receptors yielded crosslinking fractions above 80% in array-competent cells (Table S2), we conclude that all receptors in core signaling units have comparable structures at the level of the MH bundle (Fig. S1). However, the receptor dimers converge through the flexible bundle and merge into trimers of dimers at the hairpin tip (Fig. S1). The MH1 helices in one subunit of each dimer interact at the trimer axis, but their counterparts at the trimer periphery have different partners and conceivably different functional roles (54, 75–77). Together, one receptor contacts CheW and another the P5 domain of CheA to control kinase activity. The third trimer member occupies an “outboard” position (Fig. S1), where it can contact a member of the hexameric CheW rings (77) that contribute to array cooperativity, but are not essential for kinase activation or control (6).

Although the conformational and dynamic behaviors that control CheA activity at the Tsr hairpin tip remain a mystery, the kinase-ON receptor tip might need some dynamic behavior to enable CheA to move through its multi-step reaction cycle (17, 78–80). Conversely, a static receptor tip might “freeze” CheA activity, possibly at any of several steps in its reaction cycle. Accordingly, we suggest that MH bundles in the *a-d* packing arrangement have sufficiently stable structures to convey destabilizing forces through the flexible bundle to the hairpin tip, thereby promoting CheA activity (Fig. 7). It then follows that a dynamic, *x-da* MH bundle might have little structural influence on the flexible bundle, thus enabling the hairpin tip to adopt a more stable, static conformation that stops CheA activity (Fig. 8).

The bundle distortion at the dynamic junction in the AlphaFold 3 model of Tsr (Fig. 6) resembles one in the crystal structure of Tm14 (69). Pollard *et al*. suggested that structural asymmetry at this site, triggered by a chemoeffector stimulus or by adaptational modification changes, could be a way of propagating CheA-control signals to the receptor tip (69). It seems likely that the dynamic junction distortion in Tsr is a hallmark of the kinase-ON state, whereas the serine-induced OFF state could have little or no stable distortion of the MH bundle owing to elevated dynamic behavior of the *x-da* arrangement. Conceivably, the loss of a rigid structural connection like the dynamic junction distortion between the MH bundle and hairpin tip allows the receptor tip to adopt a more stable structure. The flexible bundle and glycine hinge (Fig. 8) probably play an important role in this transmission step (25–27). In vivo crosslinking approaches like those in the present study could help to elucidate the structural nature of stimulus signals traversing this region of the Tsr molecule.

## MATERIALS AND METHODS

### Bacterial Strains and Plasmids

Strains used in this study were derivatives of *E. coli* K12 strain RP437 (81). All carried extensive in-frame deletions of the MCP-family chemoreceptor genes (tsr, tar, tap, trg, aer) and five auxotrophic mutations (his, leu, met, thi, thr). Additional properties relevant to the study were: UU2610 (CheRB^-^) (82); UU2612 (CheRB^+^) (82); UU2567 (CheRBYZ^-^) (62).

Plasmid derivatives of pACYC184 (83) used in the study were: vector pKG116 (84) which confers chloramphenicol resistance and has a sodium salicylate-inducible cloning site; FRET reporter plasmid pRZ30, a pKG116 derivative that expresses *cheY-yfp* and *cheZ-cfp* under salicylate control (62); and pPA114, a pKG116 derivative that expresses wild-type *tsr* under salicylate control (47). Plasmid derivatives of pBR322 (85) used in the study were pRR48, which confers ampicillin resistance and has an isopropyl-β-D-thiogalactopyranoside (IPTG)-inducible cloning site (49), and pRR53, a pRR48 derivative that expresses wild-type *tsr* under IPTG control (49).

### Site-Directed Mutagenesis

Mutations were created in pRR53 and pPA114 by QuickChange PCR mutagenesis (86) and confirmed by sequencing.

### Growth Media

Liquid bacterial cultures were grown in T broth (1% tryptone and 0.5% NaCl wt/vol) or L broth (T broth plus 5 g/L yeast extract). Transformations were plated on L plates (1% tryptone, 0.5% NaCl wt/vol, 5 g/L yeast extract, 15% agar) containing 100 μg/mL ampicillin. Chemotaxis assays were performed on T swim plates (T broth plus 0.25% agar) containing 100 μM IPTG to induce Tsr proteins to native levels and 50 μg/mL ampicillin.

### Soft Agar Chemotaxis Assays

UU2612 transformant colonies carrying mutant pRR53 derivatives were transferred with toothpicks to T swim plates and incubated at 32.5°C for 6-7 hours.

### Quantifying Intracellular Levels of Mutant Tsr Proteins

Protein expression assays were performed in strain UU2610, as previously described (22).

### BMOE Crosslinking Assays

BMOE crosslinking assays were performed as previously described (16). Briefly, strain UU2610 carrying pRR53 and/or pPA114 CYS-reporter derivatives were grown at 30°C in tryptone broth with appropriate antibiotic and inducer concentrations. Cells were collected at mid-exponential phase by centrifugation, washed once, then resuspended in tethering buffer (87). Cell samples with or without 10 mM serine were incubated for 20 min. at 30°C before addition of BMOE (Thermo Scientific) to a final concentration of 20-200 µM. Reactions were incubated at 30°C and quenched at various times by addition of NEM (N-ethylmaleimide) to a final concentration of 10 mM. Samples were pelleted and resuspended in 2X Laemmli sample buffer (88). Samples to be reduced were resuspended in 2X Laemmli sample buffer containing 175 mM DTT (dithothreitol). Cell samples were boiled for 5-10 min. and their lysates analyzed by SDS-PAGE with 8 or 9% acrylamide gels and Tsr bands visualized by immunoblotting with polyclonal rabbit antiserum.

### In Vivo FRET-Based Kinase Assays

The experimental protocol followed (62) with hardware and data analyzed as described in (61). BMOE crosslinking FRET was performed as described (16).

### Statistics

Statistical calculations were performed in Microsoft Excel (version 16.83 for Mac) using Student’s t-test. A *p* value of <0.001 was indicated with three asterisks (***), values < 0.01 with two asterisks (**), and values <0.05 with one asterisk (*).

### Protein Structure Models

Atomic coordinates for the wild-type Tsr dimer embedded in a lipid membrane were extracted from a model of the core receptor signaling unit (54). Tsr atomic distance measurements were carried out with PyMOL (version 2.5.5 for Mac) (Schrödinger software).

### AlphaFold 3 structural predictions

The wild-type Tsr amino acid sequence was submitted to the AlphaFold 3 (beta) server (67) with molecule type set to “Protein” and copies set to “2”. The resulting fourth recycle (fold_x_model_4.cif) was converted to a *.pdb* file using PyMol (version 3.0.3 for Mac) (Schrödinger software). Crick angle deviations were determined using SamCC Turbo (68) by manually defining the cytoplasmic bundle as N-helix residues 266-389 and C-helix residues 393-516. The assignment of *x-da* packing was made based on a Crick angle of 26°±5 (89)

## ACKNOWLEDGEMENTS

Thanks to Brian Crane, Ady Vaknin, and Claudia Studdert for constructive comments on earlier drafts of the manuscript. This work was supported by research grant GM19559 (J.S.P.) from the National Institute of General Medical Sciences and the Scott A. Lloyd Memorial Graduate Fellowship at the University of Utah (G.I.R.). The Protein-DNA Core Facility at the University of Utah receives support from National Cancer Institute grant CA42014 to the Huntsman Cancer Institute.

## Author Contributions

GIR, CEF and JSP designed the experiments; GIR and CEF performed and analyzed the experiments; GIR, CEF and JSP wrote the manuscript.

## Abbreviations used

BMOE: bismaleimidoethane
CFP: cyan fluorescent protein
YFP: yellow fluorescent protein
CuPhen: Cu^2+^(1,10 phenanthroline)
CYS: cysteine
SER: serine
DTT: dithothreitol
FRET: Förster resonance energy transfer
HAMP: histidine kinases/adenylate cyclases/MCPs/phosphatases
IPTG: isopropyl-β-D-thiogalactopyranoside
MCP: methyl-accepting chemotaxis protein
MH: methylation helix
SDS-PAGE: sodium dodecyl sulfate polyacrylamide gel electrophoresis
WT or wt: wild type

