## Supplementary material for "The Structural Logic of Dynamic Signaling in the *Escherichia coli* Serine Chemoreceptor": SI materials

John S. Parkinson

This file includes:

- Figures S1 - S8
- Tables S1-S2
- SI references

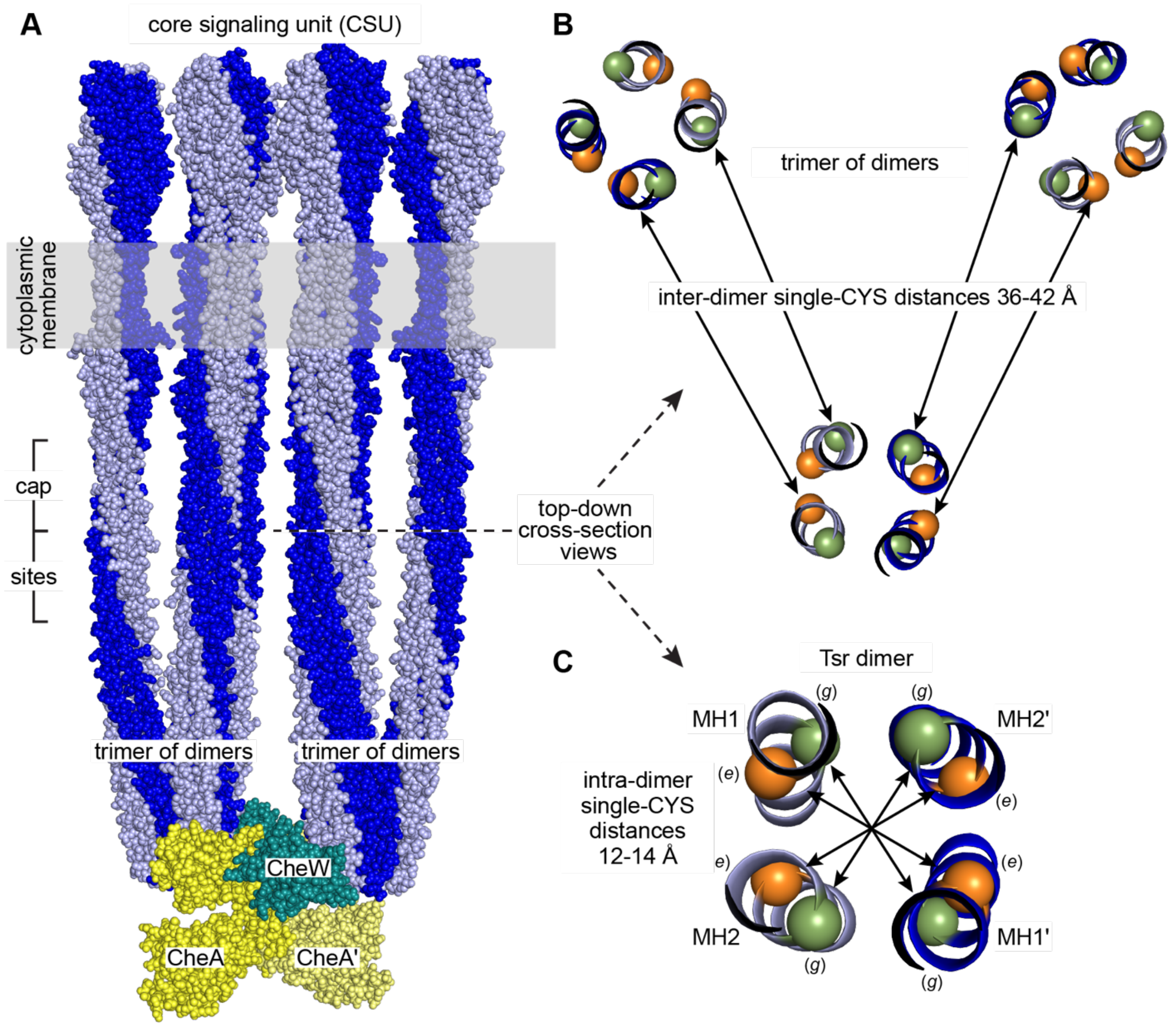


**Fig. S1.** Distances between single-cysteine (CYS) reporter sites in Tsr dimers and trimers of dimers. (A) Membrane-embedded Tsr core signaling unit (CSU); atomic coordinates from PDB: 8C5V (1). The CSU contains six Tsr dimers (light and dark blue subunits) organized in two trimers of dimers, one CheA dimer and two CheW proteins, one of which is largely hidden behind the receptors.
(B) Top-down cross-section of a Tsr trimer showing one-heptad slabs of the three dimers at the cap-sites junction of the MH bundle. Spheres are alpha-carbon atoms of the *e* (orange) and *g* (green) edge residues that flank the packing faces of the 4-helix bundles. Closest inter-dimer distances between single-CYS reporter sites were determined with PyMol 2.5.5 (Shrödinger software). (Note that the distances between the beta-carbons of cysteine residues at those reporter sites would be several Å less.)
(C) Top-down cross-section view of the reporter sites in an individual receptor dimer. Distances between the single-CYS sites within the dimer were measured with PyMol. (Note that the distances between the beta-carbons of cysteine residues at the reporter sites would be several Å less.)

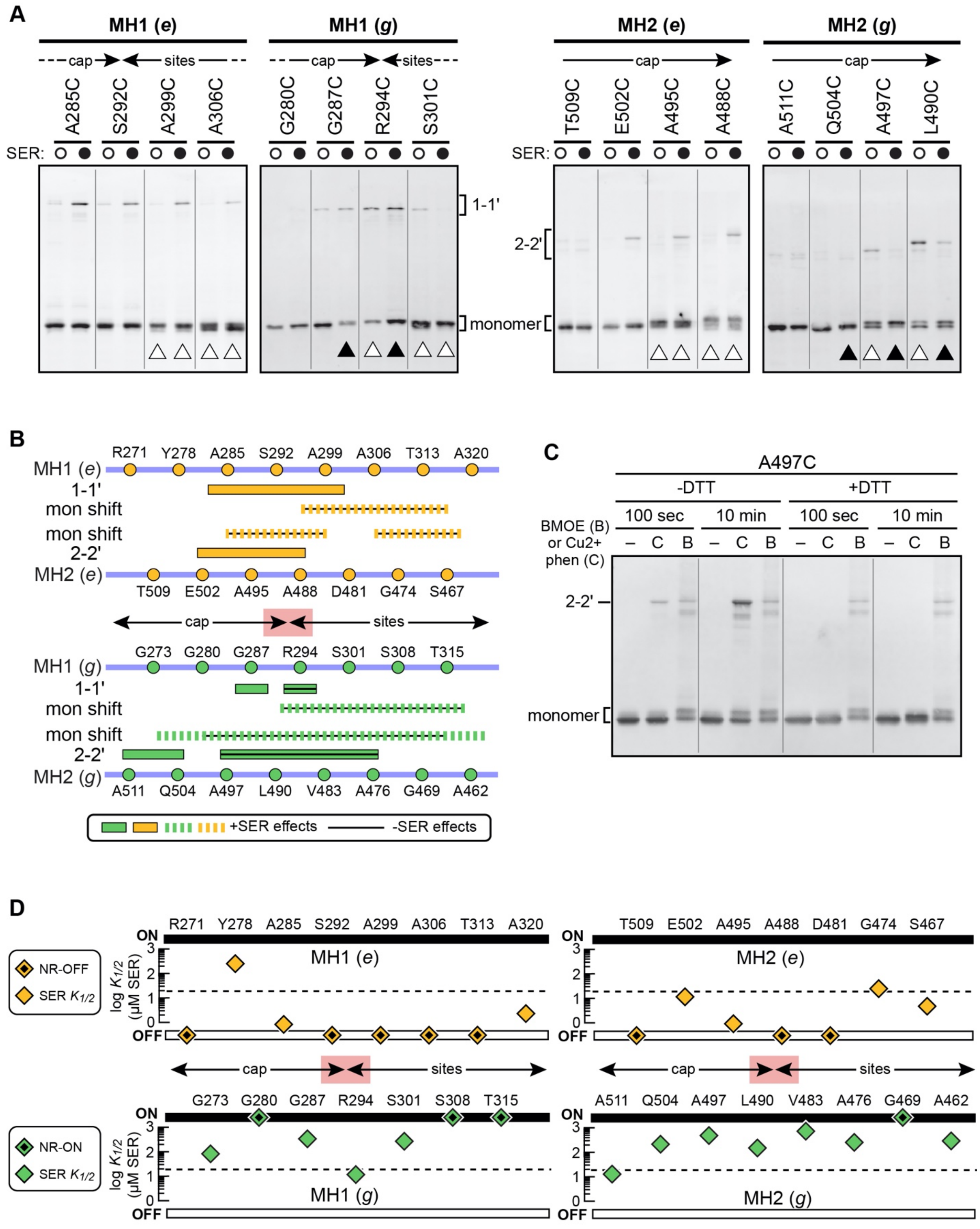


**Fig. S2.** Tsr single-CYS crosslinking data. *Legend is on following page.*

**Fig. S2.** Tsr single-CYS crosslinking data.

(A) Examples of single-CYS crosslinking gels. UU2610 cells carrying single-CYS reporter plasmids in the absence (white circle) or presence (black circle) of 10 mM serine were treated with 200 μM BMOE for 100 seconds at 30°C. Triangles indicate reporters that produced shifted monomer bands. Black triangles indicate reporters that produced a greater fraction of shifted subunits in the presence of serine. All panels show the MH bundle regions where adjacent reporters transitioned from no monomer shifts to readily apparent monomer shifts.

(B) Summary of MH bundle dynamic behaviors inferred from single-CYS reporter experiments. Lines and boxes indicate single-CYS reporters with crosslinking yields of 15% or more in the absence (lines) and/or presence (boxes) of 10 mM serine (from Fig. 2*B*). Broken lines indicate regions in which reporters produced shifted monomer bands in the presence of serine; black lines indicate band shift effects seen in the absence of serine. The red rectangle at the cap-sites border is the dynamic junction defined in Fig. 2.

(C) Modifications of A497C monomers induced by BMOE and Cu2+ phenanthroline. UU2610 cells carrying a Tsr-A497C derivative of plasmid pRR53 were treated at 30°C with no crosslinker (–), 200 μM BMOE or 300 μM Cu2+ phenanthroline for either 100 seconds or 10 minutes. All cells were pre-treated with 10 mM serine to enhance monomer bandshifts. Sample aliquots were treated with 175 mM dithiothreitol (+DTT) to reduce disulfide bonds before SDS-PAGE analysis.

(D) FRET kinase control properties of uncrosslinked Tsr single-CYS reporters. UU2567 cells carrying Tsr expression plasmid pRR53 single-CYS derivatives were tested for serine responses in FRET kinase assays (see Methods for experimental details). NR-OFF receptors exhibited no response to serine stimuli up to 10 mM and no FRET change in response to 3 mM KCN, indicative of no CheA activity. NR-ON receptors exhibited kinase activity upon KCN challenge but failed to inhibit that activity when challenged with 10 mM serine. The dashed horizontal lines indicate the $K_{1/2}$ serine response value for wild-type Tsr. The red rectangle at the cap-sites border is the dynamic junction defined in Fig. 2.

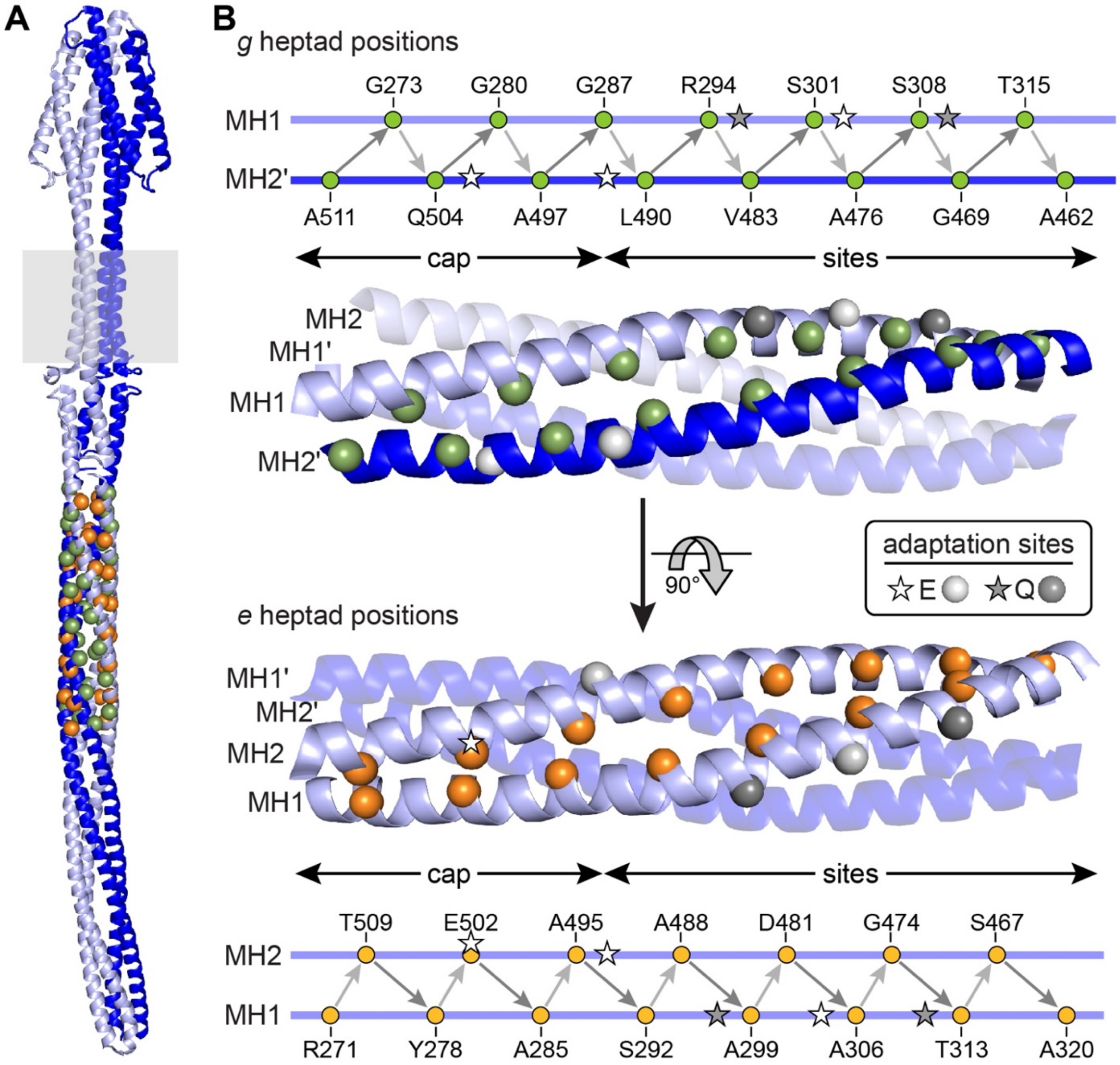


**Fig. S3.** Structural arrangements of cysteine reporter sites at bundle edge residues.
(A) A full-length membrane-embedded Tsr dimer; atomic coordinates were extracted from PDB: 8C5V (1) and analyzed with PyMol 2.5.5 (Shrödinger software). The two subunits are in different shades of blue and the membrane is depicted by a gray rectangle. The alpha-carbons of the MH bundle *e* (orange) and *g* (green) edge residues are indicated by spheres, which are shown at 1.5X scale to enhance their visibility.
(B) The positions of cysteine reporter sites are shown on the helix backbone structures and labeled in the accompanying cartoons (in which the relative positions of reporter sites are only approximate). Only one set of reporter site atoms is shown for the dimer, with the two helices in back dimmed. Distances between alpha carbons of adjacent reporter sites were measured with PyMol 2.5.5 (Schrödinger software) for the two CYS-pairs at each position in the dimer. (Note that the distances between beta-carbons of cysteine residues at the reporter sites would be several Å less.) Distances between adjacent *g*-CYS sites in the MH2'-MH1 direction (dark gray arrows) had a composite average and standard deviation of 10.2 ± 0.7 Å; corresponding values for sites in the MH1-MH2' direction (light gray arrows) were 6.6 ± 0.5 Å. Distances between *e*-CYS sites in the MH1-MH2 direction (light gray arrows) averaged 6.3 ± 0.9 Å; distances between sites in the MH2-MH1 direction (dark gray arrows) were 9.4 ± 0.6 Å.

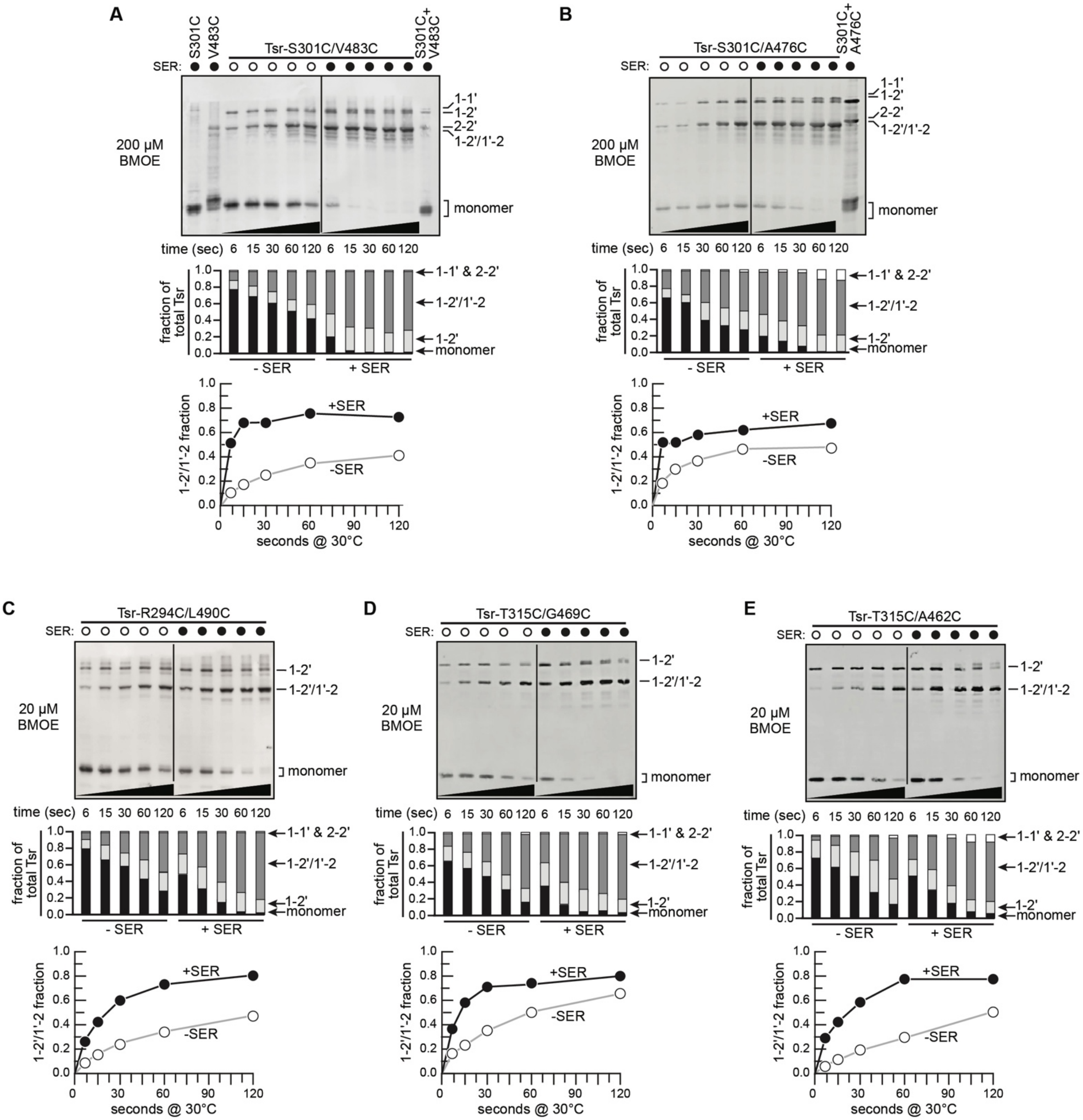


**Fig. S4.** Examples of Tsr *g/g* crosslinking timecourses. Reactions were done in cells with (black circles) and without (white circles) a 10 mM serine pretreatment. Stacked histograms show quantification of each crosslink product in the timecourse gels above them and line graphs below highlight the 1-2'/1'-2 doubly crosslinked species ±SER.

(A and B) Crosslinking reactions with 200 µM BMOE.

(C-E) Crosslinking reactions with 20 µM BMOE.

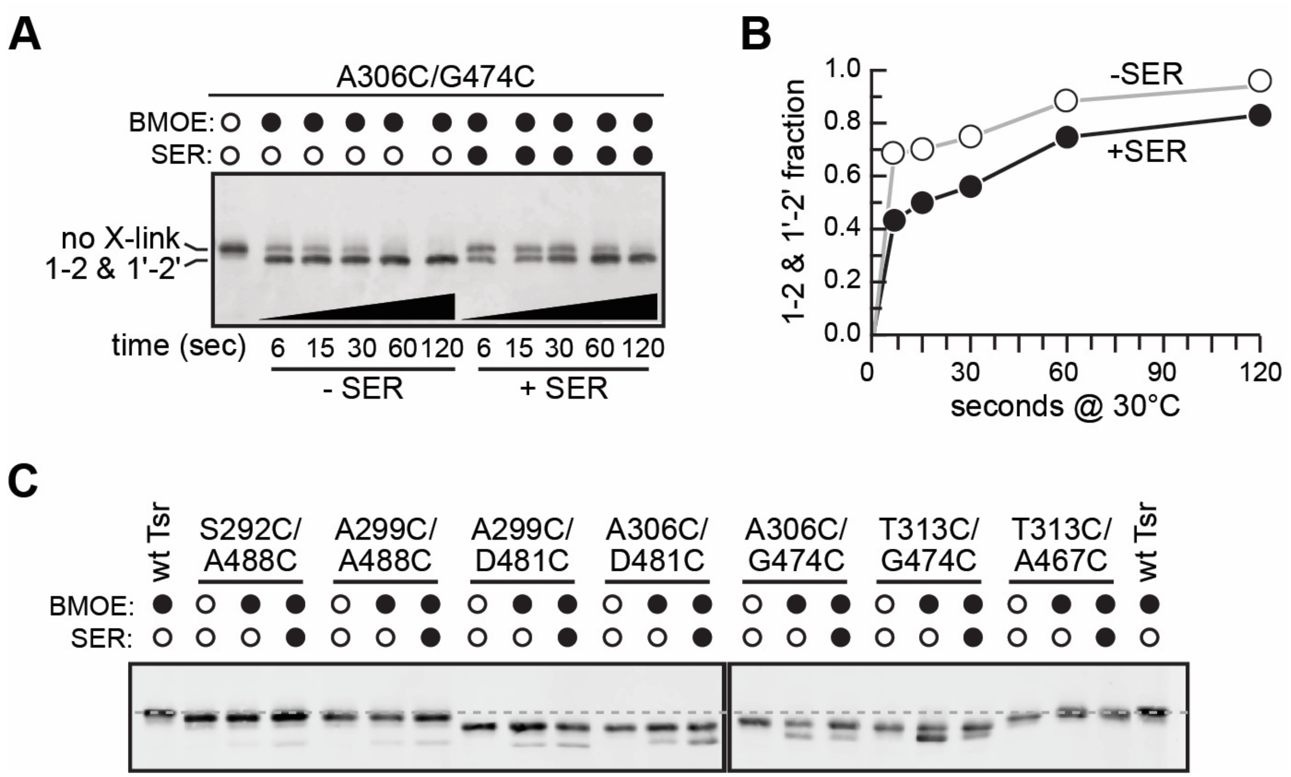


**Fig. S5.** Examples of Tsr *e/e* crosslinking. BMOE reactions were done in cells with (black circles) and without (white circles) a 10 mM serine pretreatment.

(A) Crosslinking timecourse of *e/e*-CYS reporter A306C/G474C ±SER with 50 μM BMOE.

(B) The quantified timecourse data of (A) plotted as a graph.

(C) Tsr *e/e*-CYS reporters spanning the MH sites region. Samples were incubated with or without 20 μM BMOE for 10 sec. The dashed line indicates the band position of wild-type Tsr subunits. Note that CYS-pair receptors containing the D481C or G474C reporters in MH2 have intrinsically faster mobility than the wild-type control. Note also that the relative positions of crosslinked and uncrosslinked subunits for each reporter pair depend on the distance of the reporter sites from the hairpin tip.

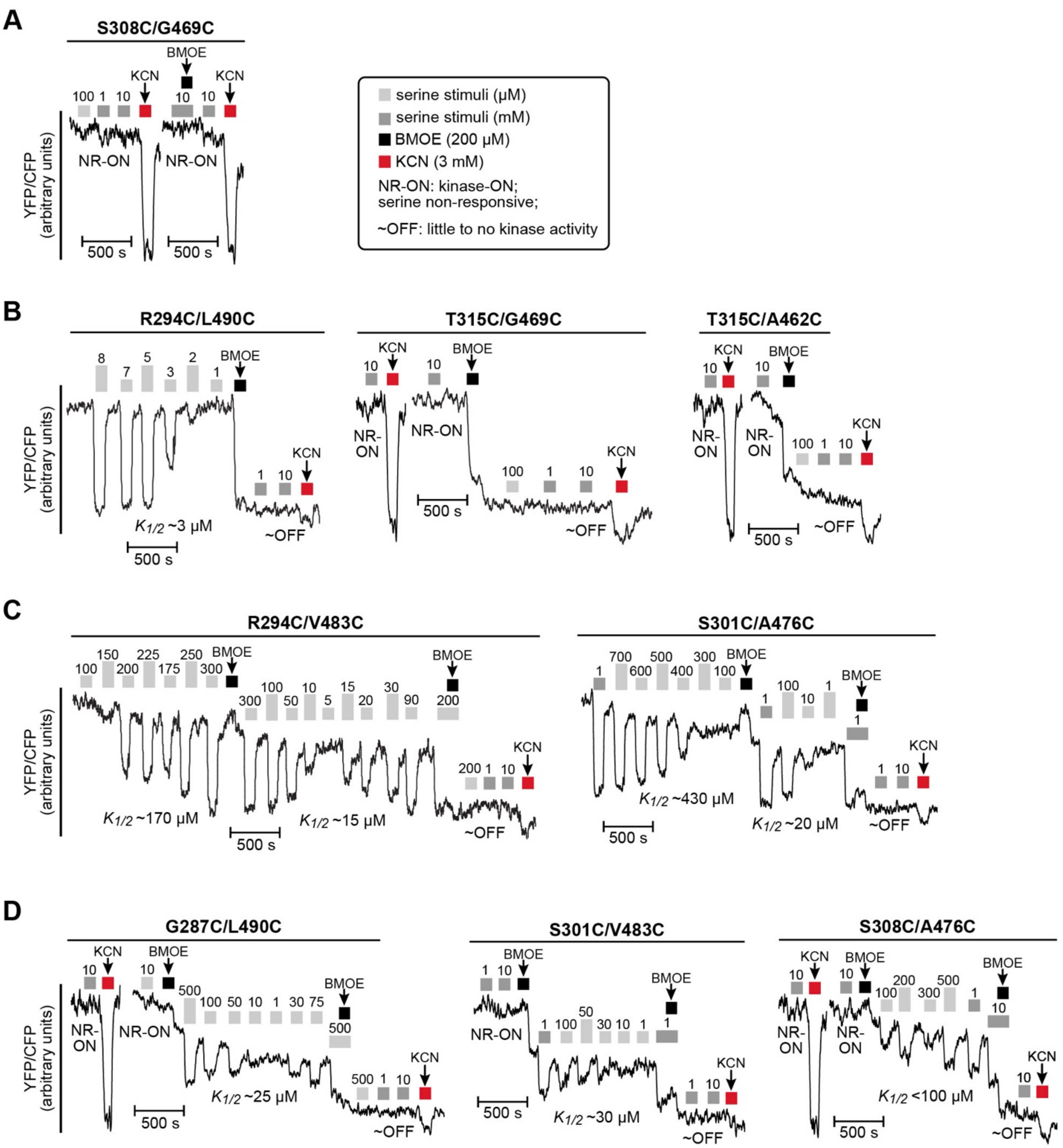


**Fig. S6.** FRET kinase assays of Tsr *g/g* reporters before and after BMOE treatment. FRET values (YFP/CFP) reflect CheA kinase activity; all plots are shown at the same scale. Small FRET drops produced by KCN treatments in the absence of CheA activity are probably due to differential effects of cellular pH changes on YFP and CFP emission efficiencies (2).

(A) Example of a serine non-responsive (NR), kinase-ON receptor; no BMOE effect.

(B) Examples of kinase-ON, serine-responsive receptors that lose kinase activity upon BMOE treatment.

(C) Examples of kinase-ON, serine-responsive receptors that become more serine-sensitive upon a first BMOE treatment, then lose all kinase activity upon a second BMOE treatment in conjunction with a saturating serine stimulus.

(D) Examples of kinase-ON, but non-responsive receptors whose kinase activity becomes serine-sensitive following an initial BMOE treatment, then becomes kinase-OFF after a second BMOE treatment in conjunction with a saturating serine stimulus.

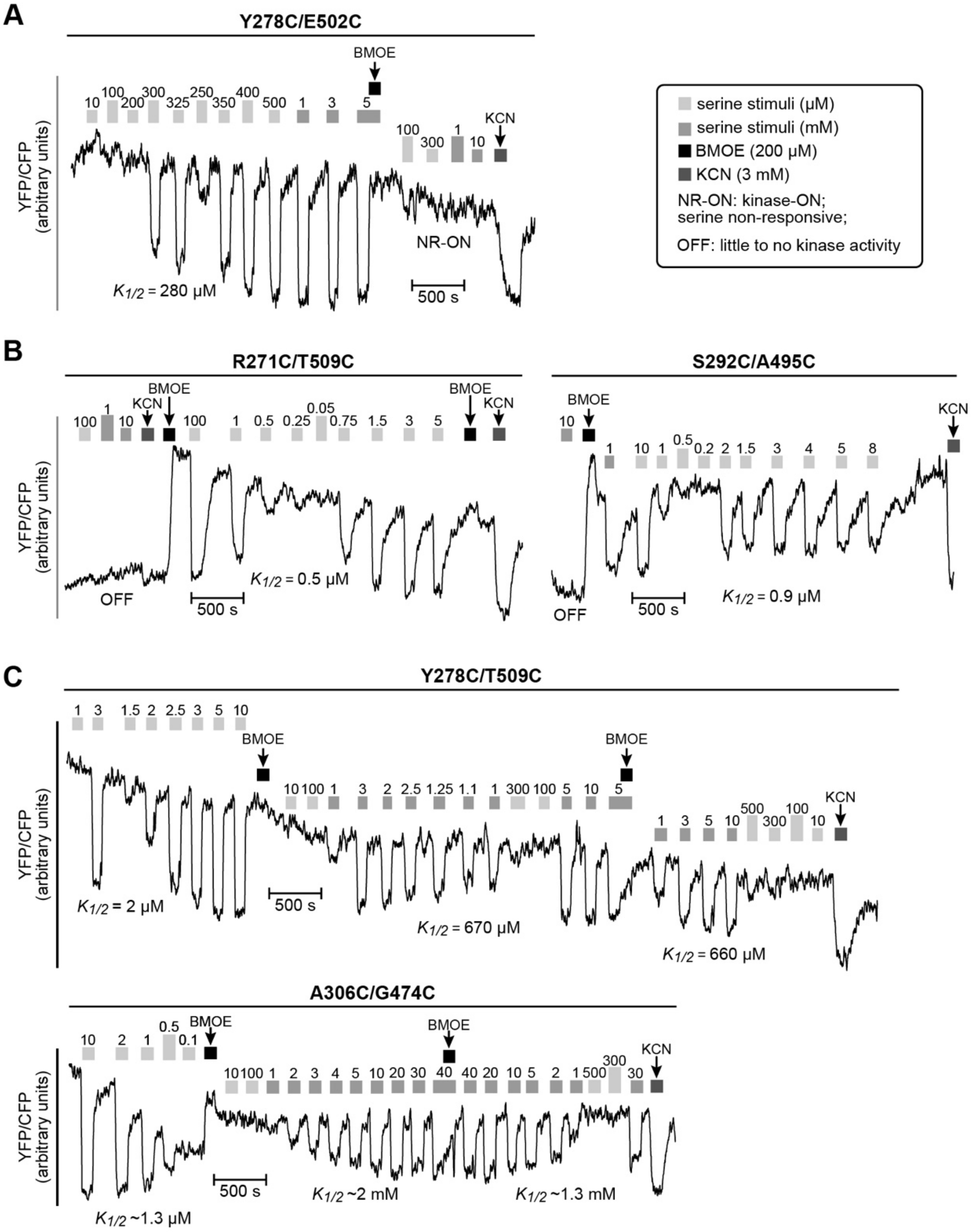


**Fig. S7.** FRET kinase assays of Tsr *e/e* reporters before and after BMOE treatment. FRET values (YFP/CFP) reflect CheA kinase activity; all plots are shown at the same scale.

(A) Example of a kinase-active, serine-responsive receptor; BMOE treatment locks kinase activity in the ON state.

(B) Examples of kinase-OFF receptors that become kinase active and serine-responsive upon BMOE treatment.

(C) Examples of kinase-active, serine-responsive receptors that become less serine-sensitive upon BMOE treatment. A second BMOE treatment in conjunction with a saturating serine stimulus produces little change in serine sensitivity. The A306C/G474C receptor lost kinase activity upon repeated serine stimuli but became kinase-active and serine-responsive again after BMOE treatment, with no further loss of kinase activity or change in serine sensitivity.

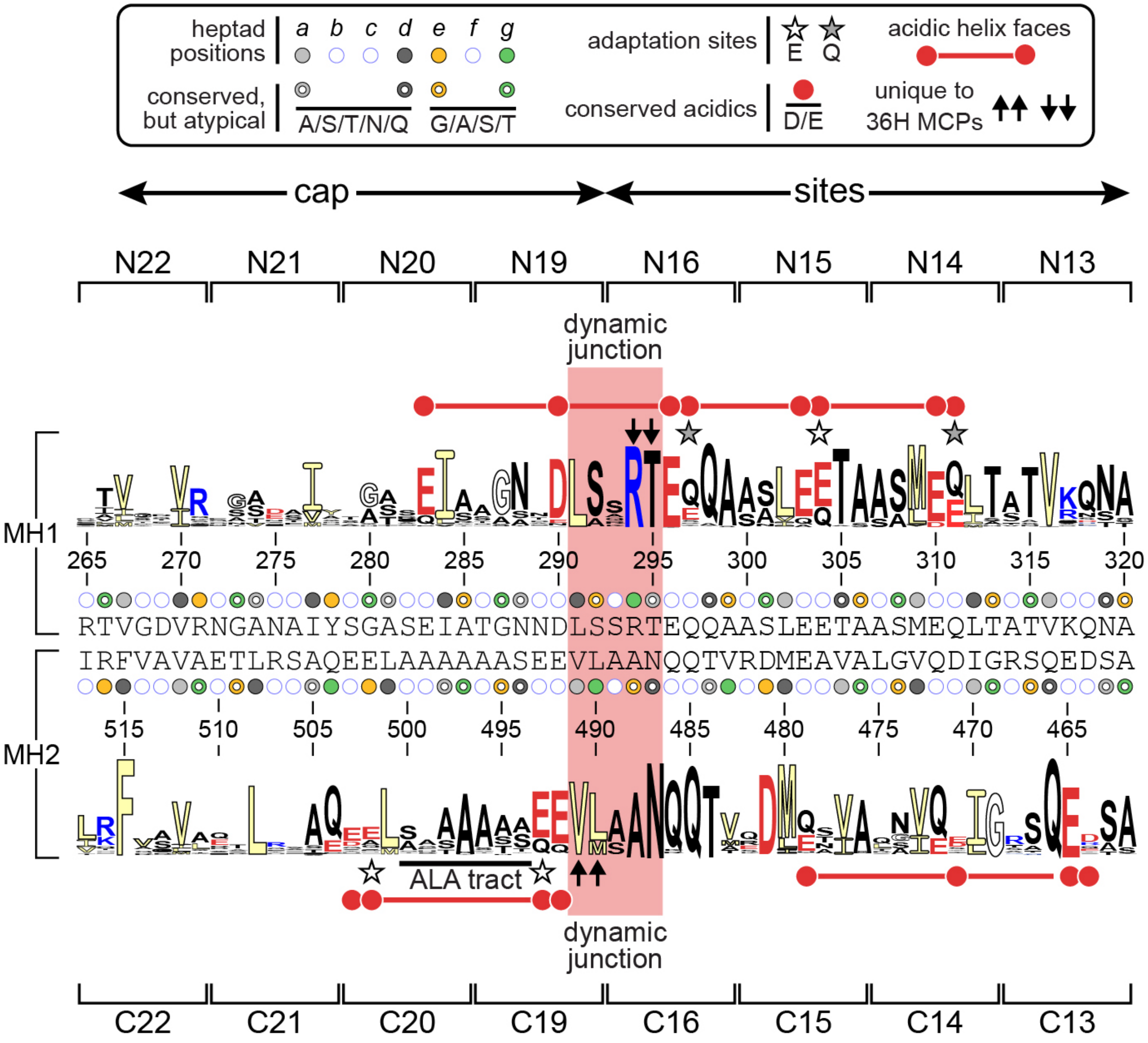


**Fig. S8.** Conserved residues and structural features of the Tsr MH bundle. The sequence logo (3) depicts the predominant residues at each MH bundle position in 2,428 nonredundant members (4) of the 36H class of chemoreceptors (5). Tsr residues and keyed heptad positions are listed between the MH1 and MH2 logos. The segment with a red background is the dynamic junction identified in this study.

**Table S1. Functional properties of Tsr CYS reporters (in pRR53 plasmid derivatives).**

| *g*-heptad reporters | | | | |
|---|---|---|---|---|
| *g*-CYS residue | function[a] | amount[b] | $K_{1/2}$ (µM SER)[c] | CheA activity[d] |
| **MH1** | | | | |
| G273C | 0.95 | 1.4 | 74 | 1.1 |
| G280C | 1.2 | 0.75 | NR-ON | 1.0 |
| G287C | 0.90 | 1.3 | 330 | 0.80 |
| R294C | 0.20 | 0.95 | 14 | 1.4 |
| S301C | 0.75 | 0.65 | 270 | 0.85 |
| S308C | 1.2 | 2.0 | NR-ON | 0.75 |
| T315C | 0.80 | 1.9 | NR-ON | 1.0 |
| **MH2** | | | | |
| A511C | 0.85 | 1.1 | 12 | 0.75 |
| Q504C | 0.90 | 1.5 | 140 | 1.3 |
| A497C | 1.0 | 1.7 | 430 | 1.2 |
| L490C | 0.85 | 0.70 | 130 | 1.4 |
| V483C | 0.90 | 1.4 | 740 | 1.4 |
| A476C | 0.60 | 0.80 | 240 | 0.85 |
| G469C | 0.15 | 0.50 | NR-ON | 0.85 |
| A462C | 0.45 | 1.5 | 250 | 1.2 |

| *g/g* reporter pairs | | | | |
|---|---|---|---|---|
| *g/g* CYS residues | function[a] | amount[b] | $K_{1/2}$ (µM SER)[c] | CheA activity[d] |
| **MH1/MH2** | | | | |
| G273C/A511C | 0.60 | 0.75 | 97 | 0.90 |
| G273C/Q504C | 0.95 | 0.70 | 410 | 1.2 |
| G280C/Q504C | 0.55 | 0.75 | NR-ON | 1.6 |
| G280C/A497C | 0.15 | 0.80 | NR-ON | 1.1 |
| G287C/A497C | 0.50 | 1.2 | 347.10 | 0.70 |
| G287C/L490C | 0.55 | 1.1 | NR-ON | 0.75 |
| R294C/L490C | 0.20 | 0.65 | 3.0 | 1.6 |
| R294C/V483C | 0.30 | 0.80 | 170 | 1.4 |
| S301C/V483C | 0.80 | 1.2 | NR-ON | 0.50 |
| S301C/A476C | 0.25 | 0.60 | 480 | 0.60 |
| S308C/A476C | 0.20 | 0.50 | NR-ON | 1.2 |
| S308C/G469C | 0.15 | 0.85 | NR-ON | 1.3 |
| T315C/G469C | 0.15 | 0.65 | NR-ON | 1.3 |
| T315C/A462C | 0.50 | 1.6 | NR-ON | 1.3 |

| *e*-heptad reporters | | | | |
|---|---|---|---|---|
| *e*-CYS residue | function[a] | amount[b] | K1/2 (µM SER)[c] | CheA activity[d] |
| **MH1** | | | | |
| R271C | 0.10 | 0.85 | NR-OFF | 0.00 |
| Y278C | 0.65 | 0.85 | 310 | 0.85 |
| A285C | 0.45 | 1.2 | 0.90 | 1.3 |
| S292C | 0.20 | 0.85 | NR-OFF | 0.00 |
| A299C | 0.20 | 0.70 | NR-OFF | 0.00 |
| A306C | 0.30 | 0.60 | NR-OFF | 0.00 |
| T313C | 0.30 | 0.65 | NR-OFF | 0.00 |
| A320C | 0.10 | 0.65 | 2.1 | 1.3 |
| **MH2** | | | | |
| T509C | 0.10 | 1.1 | NR-OFF | 0.00 |
| E502C | 0.65 | 1.1 | 16 | 1.3 |
| A495C | 0.60 | 1.9 | NR-OFF | 0.75 |
| A488C | 0.35 | 0.65 | NR-OFF | 0.00 |
| D481C | 0.10 | 0.75 | NR-OFF | 0.00 |
| G474C | 0.65 | 0.75 | 29 | 1.3 |
| S467C | 0.15 | 0.90 | 5.0 | 1.6 |

| *e/e* reporter pairs | | | | |
|---|---|---|---|---|
| *e/e* CYS residues | function[a] | amount[b] | K1/2 (µM SER)[c] | CheA activity[d] |
| **MH1/MH2** | | | | |
| R271C/T509C | 0.05 | 0.75 | NR-OFF | 0.00 |
| Y278C/T509C | 0.35 | 1.3 | NR-OFF | 1.2 |
| Y278C/E502C | 0.75 | 1.1 | 2.0 | 1.1 |
| A285C/E502C | 0.50 | 1.1 | 280 | 1.2 |
| A285C/A495C | 0.15 | 0.75 | 84 | 0.00 |
| S292C/A495C | 0.10 | 2.2 | NR-OFF | 0.00 |
| S292C/A488C | 0.10 | 1.8 | NR-OFF | 0.00 |
| A299C/A488C | 0.20 | 1.5 | NR-OFF | 0.00 |
| A299C/D481C | 0.10 | 2.6 | NR-OFF | 0.00 |
| A306C/D481C | 0.10 | 1.6 | NR-OFF | 0.00 |
| A306C/G474C | 0.50 | 1.0 | 1.4 | 0.85 |
| T313C/G474C | 0.65 | 1.2 | 3.4 | 1.8 |
| T313C/S467C | 0.15 | 2.1 | 1.0 | 1.3 |
| A320C/S467C | 0.70 | 1.5 | 18 | 1.4 |

[a] Colony size relative to wild-type Tsr control on tryptone soft agar; pRR53 mutant plasmids in strain UU2612.

[b] Amount of mutant protein relative to wild-type Tsr control; pRR53 mutant plasmids in strain UU2610.

[c] FRET assay of pRR53 mutant plasmids in strain UU2567; NR-ON: activity, no SER response; NR-OFF: no activity, no SER response.

[d] Activity relative to wild-type Tsr control in FRET assay of pRR53 mutant plasmids in strain UU2567.

All data rounded as follows: values <1 rounded to 0.05; values <10 rounded to 0.1; values <100 rounded to 1; values <1000 rounded to 10.

**Table S2. Raw crosslinking data for Tsr CYS-pair reporters**

| 1-2'/1'-2 fraction | | | | |
|---|---|---|---|---|
| *g/g*-CYS pairs | -SER | +SER | N | p value |
| G273C/A511C | 0.08 ± 0.02 | 0.37 ± 0.02 | 3 | <0.001 |
| G273C/Q504C | 0.18 ± 0.02 | 0.70 ± 0.02 | 3 | <0.001 |
| G280C/Q504C | 0.17 ± 0.02 | 0.78 ± 0.05 | 3 | <0.001 |
| G280C/A497C | 0.17 ± 0.02 | 0.43 ± 0.02 | 3 | <0.001 |
| G287C/A497C | 0.20 ± 0.02 | 0.55 ± 0.03 | 3 | <0.001 |
| G287C/L490C | 0.34 ± 0.02 | 0.78 ± 0.02 | 5 | <0.001 |
| R294C/L490C | 0.69 ± 0.09 | 0.70 ± 0.12 | 5 | 0.89 |
| R294C/V483C | 0.31 ± 0.04 | 0.67 ± 0.03 | 5 | <0.001 |
| S301C/V483C | 0.55 ± 0.03 | 0.72 ± 0.03 | 5 | 0.002 |
| S301C/A476C | 0.43 ± 0.03 | 0.74 ± 0.03 | 5 | <0.001 |
| S308C/A476C | 0.23 ± 0.04 | 0.74 ± 0.14 | 5 | 0.004 |
| S308C/G469C | 0.12 ± 0.04 | 0.53 ± 0.06 | 5 | <0.001 |
| T315C/G469C | 0.82 ± 0.05 | 0.70 ± 0.13 | 5 | 0.21 |
| T315C/A462C | 0.83 ± 0.02 | 0.82 ± 0.04 | 5 | 0.66 |

| 1-2 + 1'-2' fraction | | | | |
|---|---|---|---|---|
| *e/e*-CYS pairs | -SER | +SER | N | p value |
| R271C/T509C | 0.15 ± 0.03 | 0.18 ± 0.05 | 3 | 0.40 |
| Y278C/T509C | 0.50 ± 0.11 | 0.28 ± 0.04 | 3 | 0.03 |
| Y278C/E502C | 0.60 ± 0.02 | 0.68 ± 0.11 | 3 | 0.27 |
| A285C/E502C | 0.46 ± 0.04 | 0.26 ± 0.02 | 3 | 0.001 |
| A285C/A495C | 0.16 ± 0.03 | 0.04 ± 0.01 | 3 | 0.002 |
| S292C/A495C | 0.18 ± 0.01 | 0.18 ± 0.03 | 3 | 0.74 |
| S292C/A488C | 0.06 ± 0.01 | 0.08 ± 0.01 | 3 | 0.015 |
| A299C/A488C | 0.06 ± 0.02 | 0.09 ± 0.03 | 3 | 0.24 |
| A299C/D481C | 0.15 ± 0.06 | 0.20 ± 0.01 | 3 | 0.21 |
| A306C/D481C | 0.21 ± 0.01 | 0.26 ± 0.03 | 3 | 0.03 |
| A306C/G474C | 0.32 ± 0.03 | 0.21 ± 0.05 | 3 | 0.03 |
| T313C/G474C | 0.43 ± 0.04 | 0.26 ± 0.07 | 3 | 0.02 |
| T313C/S467C | 0.53 ± 0.06 | 0.41 ± 0.06 | 3 | 0.08 |
| A320C/S467C | 0.23 ± 0.01 | 0.11 ± 0.03 | 3 | 0.002 |

Values in the -SER (no serine) and +SER (serine pre-treatment) columns are means ± standard deviations for N independent experiments; *p* values were determined by Student's t-test.
